# Differential response to LSD1 inhibition in prostate cancer reveals a YAP1 vulnerability in the neuroendocrine subtype

**DOI:** 10.1101/2024.01.17.576106

**Authors:** SM Jasim, R Kumar, S Kim, TEG Krueger, IM Coleman, SL Dalrymple, L Antony, DM Rosen, B Hanratty, RA Patel, PS Nelson, Z Aghaei, JH Beumer, MC Haffner, TL Lotan, A Mandl, MA Carducci, S Kachhap, JT Isaacs, SR Denmeade, HY Rienhoff, LA Sena, WN Brennen

## Abstract

Progression to lethal metastatic castration-resistant prostate cancer (mCRPC) is driven in part by epigenetic modulators such as LSD1 (*KDM1A*), a lysine-specific demethylase. Yet, mCRPC is increasingly recognized as a highly heterogeneous disease whose classification into subtypes is defined by the extent of androgen receptor (AR) and/or neuroendocrine (NE) characteristics. Meanwhile, the role of LSD1 in driving the different subtypes of mCRPC has remained unclear. Here, we assess the necessity of LSD1 in driving progression of mCRPC subtypes including AR+/NE- (ARPC), AR-/NE+ (NEPC), AR+/NE+ (amphicrine; AMPC), and AR-/NE- (double-negative; DNPC) through the use of LSD1 inhibitors in clinical development. LSD1 inhibition (LSD1i) was observed to be highly effective in restricting growth of NEPC, and efficacy was associated with *TP53* loss-of-function. Mice bearing NEPC patient-derived xenografts treated with the LSD1 inhibitors, bomedemstat (MK-3543) or iadademstat (ORY-1001), exhibited suppression of the NE transcriptional profile, including ASCL1. LSD1i also induced expression and activity of YAP1, a non-NE transcription factor canonically silenced in NEPC (YAP^OFF^ cancer), thereby switching NEPC from a YAP^OFF^ to a YAP^ON^ cancer class. Therapeutically-induced YAP^ON^ NEPC tumors exhibited cell cycle arrest and repression of proliferative transcriptional programs. Importantly, the LSD1i-mediated YAP^ON^ state induced sensitivity to an inhibitor of YAP/TEAD function, IAG933, which extended antitumor efficacy against NEPC. Altogether, these findings indicate that patients diagnosed with NEPC may obtain greater relative benefit from LSD1-targeted therapies compared to those with other mCRPC subtypes and that dual inhibition of LSD1 and YAP/TEAD function demonstrates a promising treatment strategy potentially extending to other YAP^OFF^ cancers.

**Significance:** Across prostate cancer subtypes, NEPC is exceptionally responsive to LSD1 inhibition and this response is enhanced in combination with a YAP/TEAD disruptor which may improve patient selection and outcomes.

**Graphical abstract:** 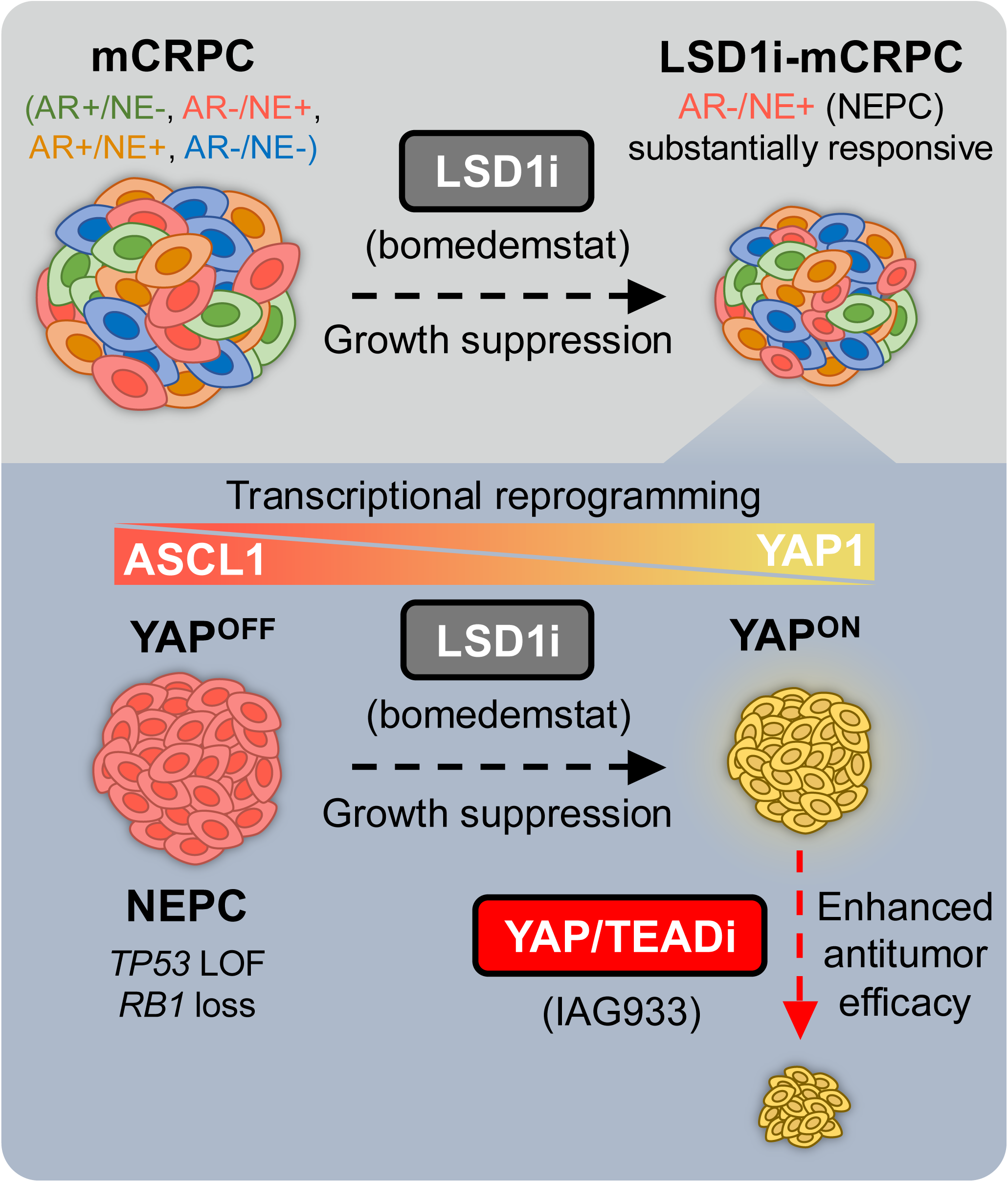

## Introduction

Despite significant advancements in therapy, prostate cancer persists as a leading cause of cancer-related death in men worldwide. These deaths are mainly attributable to progression of metastatic disease to castration-resistant prostate cancer (mCRPC) (1). Most mCRPC is driven by androgen receptor (AR) signaling and classified as adenocarcinoma (ARPC). However, as a consequence of mCRPC tumor cell heterogeneity, treatment pressure from AR signaling inhibitors leads to the emergence of alternative prostate cancer subtypes. Up to 20% of prostate cancer tumors demonstrate loss of dependence on AR signaling and acquisition of neuroendocrine (NE) features (NEPC) (2). NEPC is an aggressive mCRPC subtype without effective treatments and exhibits a median survival of less than two years (3).

Genomic alterations in *PTEN*, *RB1*, and *TP53* predispose mCRPC tumor cells to undergo transdifferentiation from ARPC to NEPC (4,5). This transformation proceeds through distinct transcriptional states. During early transition initiation, stress-responsive transcription factors NFkB, JUN, and YAP1 are activated (6). Subsequently, intermediate reprogramming factors within POU and SOX families become highly enriched. In late transformation, neuronal transcription factors NEUROD1 and ASCL1 establish a NE transcriptional profile (6). These transcriptional states are also observed in other treatment-induced NE carcinomas such as small cell lung cancer (SCLC) which is defined by ASCL1, NEUROD1, POU2F3, or YAP1 subtypes (7).

In addition to genomic and transcriptional alterations, mCRPC tumor cells undergoing transdifferentiation also manifest marked changes in their epigenetic landscape. Histone modifications, for example, differ significantly between ARPC and NEPC (8). In particular, the histone modifier lysine-specific demethylase 1 (LSD1 or *KDM1A*), is highly expressed in NEPC tumors compared to ARPC and correlates with worse prognosis (9). Small molecule inhibitors of LSD1 are in late-stage clinical development for cancer and other diseases (10). Antitumor efficacy from LSD1 inhibition (LSD1i) in animal studies appears promising in NEPC models expressing wildtype *TP53* (9). However, nearly 70% of NEPC patients exhibit a *TP53* alteration, which contributes to the overall heterogeneity, plasticity, and lethality of the disease (5,8,11). Efficacy of LSD1i in *TP53*-deficient NEPC remains unknown. Furthermore, well-characterized response rates from advanced prostate cancer models treated with LSD1i *in vivo* are scarce. Addressing these two gaps in the field will guide LSD1i use in a stratified patient population to potentially improve outcomes as these therapies complete late-stage clinical trials.

Here, we determined that antitumor efficacy by LSD1i in mCRPC models was associated with *TP53* LOF. The most clinically aggressive subtype, NEPC, showed the highest degree of growth suppression by LSD1i. Transcriptionally, LSD1i suppressed ASCL1 expression, indicating a role for LSD1 as an upstream epigenetic driver of the NE transcriptional program in NEPC tumors. Furthermore, YAP1, which is canonically silenced in NEPC (12), was de-repressed, suggesting LSD1 maintains NEPC as a YAP^OFF^ cancer class through epigenetic regulation. LSD1i-mediated YAP1 upregulation switched NEPC from a YAP^OFF^ to a YAP^ON^ state, which induced sensitivity to IAG933, a clinical-grade YAP/TEAD disruptor. Collectively, these data demonstrate that LSD1 epigenetically sustains the NE transcriptional profile and that its inhibition substantially impairs NEPC growth and dysregulates this phenotype, leading to induction of a YAP^ON^ state vulnerable to YAP1-targeted therapy.

## Results

### mCRPC response to LSD1 inhibition is most effective in NEPC and associated with *TP53* LOF

To determine which subgroups of prostate cancer patients would potentially benefit most from LSD1-targeted therapy, mice bearing cell line- and patient derived-xenografts (CDX/PDXs) across 12 different models encompassing the four major subtypes of advanced mCRPC observed clinically were treated with bomedemstat, a selective LSD1i in late-stage clinical development. We tested CDX/PDXs in male NSG mice representing the following mCRPC subtypes: AR+/NE- (ARPC; n=6 models), AR+/NE+ (amphicrine; AMPC; n=2 models), AR-/NE- (double negative; DNPC; n=2 models), and AR-/NE+ (NEPC; n=2 models). Among ARPC models, LAPC4, CWR22-RH, and VCaP exhibited the greatest reduction in tumor growth upon treatment with LSD1i (53-66%), whereas LNCaP, C4-2B, and LN-95 appeared unresponsive (Fig. 1A). Differences in response were confirmed by histology using the proliferation marker, Ki67, whose abundance was reduced in LSD1i-treated VCaP tumors but remained elevated in LSD1i-treated LNCaP tumors (Fig. S1A). In AMPC models, LSD1i dramatically reduced LvCaP-2R tumor growth but 22Rv1 was minimally responsive (Fig. 1B). In DNPC models, both DU145 and PC3 were responsive to LSD1i, though antitumor activity was more pronounced against DU145 (Fig. 1C). Meanwhile, NEPC models responded exceptionally well to LSD1i with substantial growth impairment in both NCI-H660 and LTL331R (Fig. 1D). Individual LTL331R tumor growth depicted by spider plot shows a subset of exceptional responders experiencing prolonged growth impairment with bomedemstat treatment (Fig. 1E). Tumors resumed growth upon treatment cessation, suggesting persistent LSD1i is required to maintain tumor growth control. Similar antitumor effects were achieved using a second clinically-tested small molecule inhibitor of LSD1, iadademstat (Fig. S2A, B). Moreover, LSD1i did not affect mouse body weights in any of these models, even with prolonged daily treatment for over 30 days (Fig. S3A-D).

**Figure 1.**
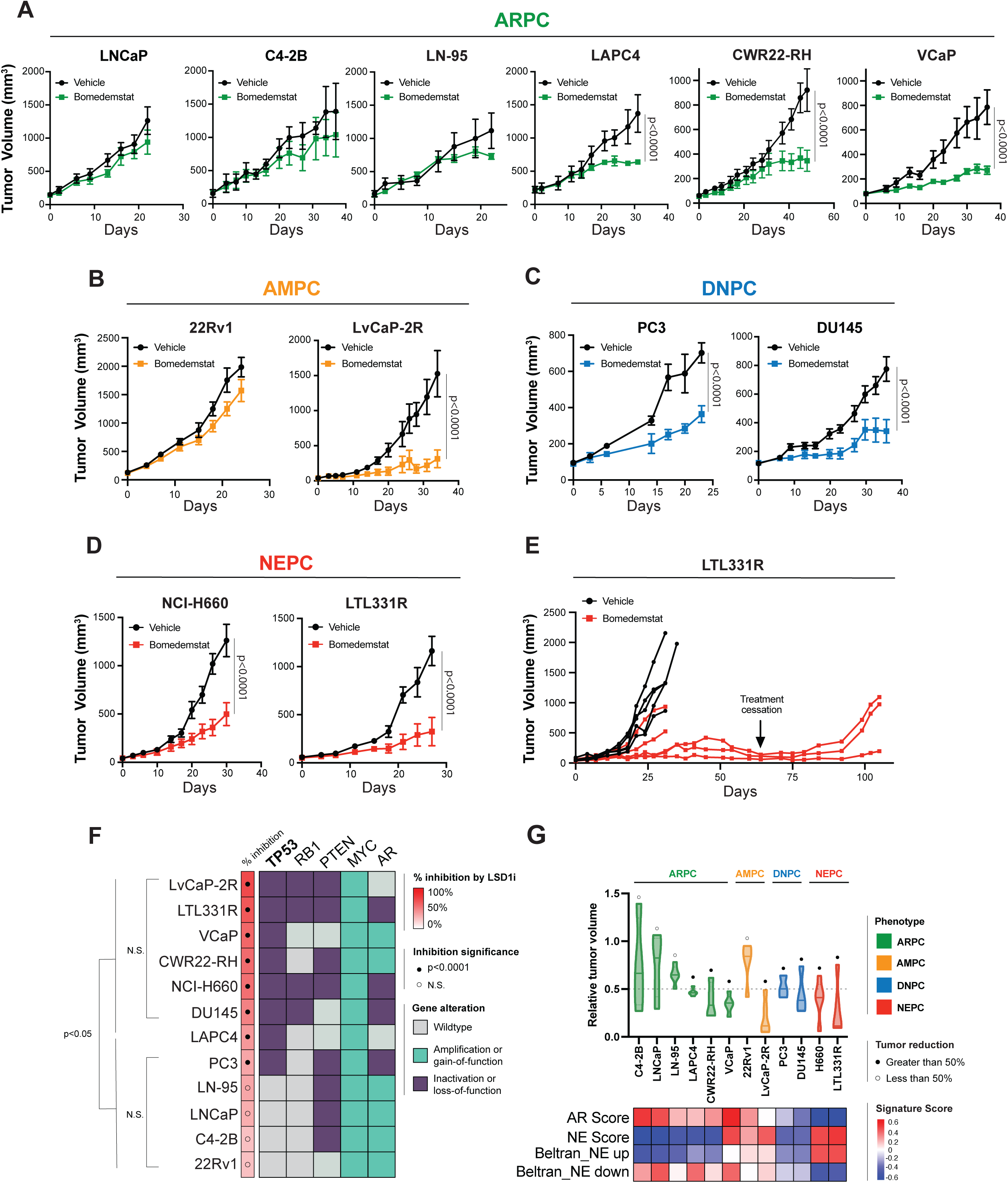
mCRPC response to LSD1i is most effective in NEPC and associated with *TP53* LOF. **A,** Tumor growth curves depicting days on treatment with response to bomedemstat (dosed PO daily at 40 mg/kg) or vehicle control in male NSG mice bearing mCRPC models of ARPC: LNCaP (n=5), C4-2B (n=3-4), LN-95 (n=4-6), LAPC4 (n=4-5), CWR22-RH (n=3-4), VCaP (n=5), **B,** AMPC: 22Rv1 (n=5), LvCaP-2R (n=5-6), **C,** DNPC: PC3 (n=3), DU145 (n=5), and **D**, NEPC: NCI-H660 (n=5), LTL331R (n=5). N=mice per condition. Data presented as the mean ±SEM; p-values determined using linear mixed model (LMM) with Šidák correction for multiple comparisons. **E,** Spider plot of individual LTL331R tumors depicting a subset of mice receiving continuous daily dosing of bomedemstat for 64 days (black arrow), when treatment was discontinued. **F,** mCRPC models ranked by greatest response to LSD1i therapy. Percent inhibition calculated from each model’s end-of-study LSD1i-treated tumor volume relative to respective vehicle-treated tumor volume mean. Inhibition significance between LSD1i-treated and respective vehicle-treated tumors determined by two-way ANOVA (or LMM) and indicated by black circle in % inhibition column. Significant differences in tumor growth between mCRPC models indicated by unpaired two-tailed *t*-test with p-value<0.05. Mutational/functional status of the most frequently perturbed genes (*TP53*, *RB1*, *PTEN*, *MYC*, and *AR*) in mCRPC depicted for all models. Gene or functional status classified using publicly-available resources. **G,** End-of-study relative tumor volumes normalized to vehicle control in each experiment. Tumor mass reduction greater than 50% indicated by black circle on plot. Subtype classification indicated by color code. AR and NE scores quantified via multi-gene signatures with GSVA scores. Data presented as the median ± SEM.

To understand the molecular determinants of differential response to LSD1i, mCRPC models were ranked according to bomedemstat efficacy calculated as percent inhibition of treatment relative to vehicle using end-of-study tumor volumes (Fig. 1F). We then queried five of the most frequently altered genes in advanced prostate cancer patients (*TP53*, *RB1*, *PTEN*, *MYC*, *AR*) (13) across the mCRPC models used here *via* previously published data (14–19) or publicly-available resources including Gene Expression Omnibus (GEO, NCBI) and DepMap dataset (DepMap 26Q1, Broad Institute). Among these genomic alterations, deficiency/loss-of-function (LOF) in *TP53* and *RB1* emerged as the most common correlates in models responsive to LSD1i therapy (Fig. 1F). Enhanced response to LSD1i in prostate cancer models with *RB1* loss is consistent with a prior report (20). However, we observed up to 65% tumor mass reduction in mice treated with bomedemstat relative to vehicle in LAPC4, CWR22-RH, and VCaP despite these models harboring functional *RB1*. Although not altered, RB1 expression is low in PC3, a model that responds to LSD1i as shown in this study (Fig. 1C) and others (21). Invariably, however, *TP53* deficiency/LOF was associated with response to bomedemstat therapy in all models, whereas *TP53* was functional in non-responders (Fig. 1F). Therefore, in these mCRPC models, the correlation between *TP53* LOF and bomedemstat sensitivity suggests a potential predictive indicator of response to LSD1i in addition to *RB1*. Between mCRPC subtypes, models of NEPC and those enriched for a NE signature had the most pronounced reductions in tumor volume following LSD1i (>60%) compared to other subtypes with differential responses summarized in Figure 1G. Clinically, it is well known that the majority of patients diagnosed with NEPC lack functional *TP53* and/or *RB1* (4,5). Given that loss of *TP53* and *RB1* is a frequent characteristic in NEPC, LSD1i therapy may be especially beneficial for patients classified with this subtype. A prior report demonstrated that LSD1i induces *TP53* activity in *TP53*-proficient NEPC models (9). However, the effects of LSD1i in clinically-relevant *TP53*-deficient and *RB1* loss NEPC models have not been previously assessed.

### LSD1 sustains the neuroendocrine transcriptional profile and ASCL1 expression in NEPC

Having observed that LSD1i response is associated with *TP53* LOF in addition to previously characterized *RB1* loss, we chose to interrogate the transcriptional consequences of targeting LSD1 in NEPC due to the frequency of *TP53* and *RB1* genetic alterations in this subtype. To understand the transcriptional effects of LSD1i in NEPC, we performed bulk RNA-seq on the NEPC transdifferentiation PDX model, LTL331R, treated with bomedemstat. Our analysis identified 10,120 differentially regulated genes between LSD1i-treated (n=5) and vehicle-control (n=3) tumors with 3,075 enriched two-fold or greater but only 996 genes significantly downregulated by two-fold or more (Fig. 2A, B). Given that LSD1 is generally associated with transcriptional repression, a global increase in gene expression was expected with bomedemstat treatment. However, among the 996 downregulated genes, those specifying the NE lineage were notably enriched. Suppression of the NE phenotype in LTL331R PDXs treated with bomedemstat was demonstrated across several signatures using gene set enrichment analysis (GSEA) (Fig. 2C, D). The NE master transcription factor *ASCL1* was also significantly downregulated as shown by RNA-seq in LTL331R PDXs (Fig. 2E) and by qPCR in NCI-H660 spheroids using iadademstat (Fig. 2F). Furthermore, ASCL1 protein abundance was also markedly attenuated by LSD1i *in vivo* using immunohistochemistry (IHC) (Fig. 2G, H) and *in vitro* using Western blot (Fig. 2I). Therefore, LSD1i leads to downregulation of the NE transcriptional profile and ASCL1 in NEPC, corroborating similar findings in SCLC using iadademstat (22,23).

**Figure 2.**
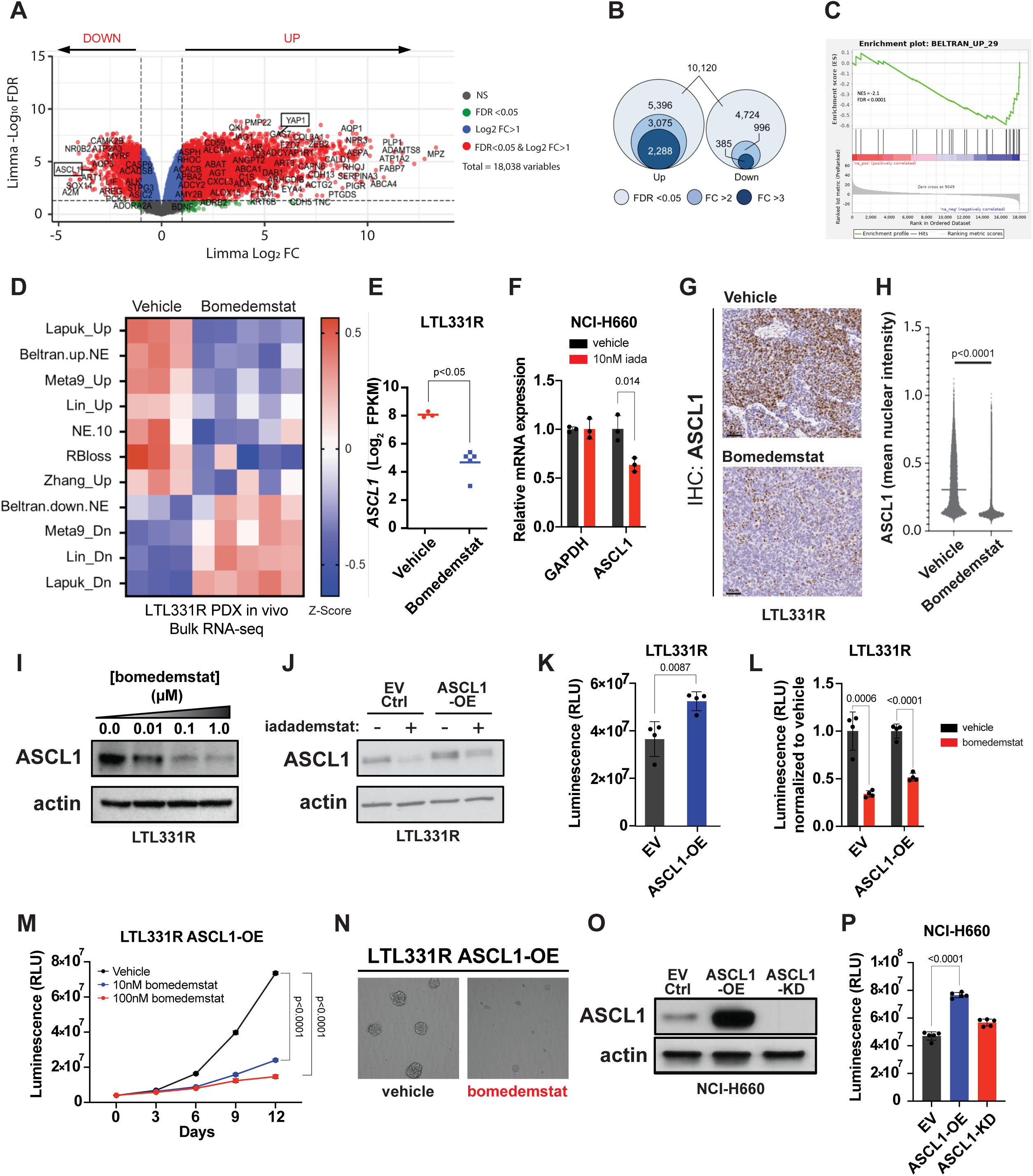
LSD1 sustains neuroendocrine transcription and ASCL1 expression in NEPC. **A,** Volcano plot from bulk RNA-seq data on LTL331R PDX end-of-study tumors treated with vehicle (n = 3) or bomedemstat (n=5). Genes significantly altered by LSD1i with FDR<0.05 and Log_2_FC>1 depicted in red. **B,** Comparison of genes significantly up- or down-regulated in bomedemstat-treated LTL331R PDX tumors compared to vehicle control. Number of genes with FDR<0.05 and FC>2 or 3 indicated. **C,** GSEA on bomedemstat-treated LTL331R PDX tumors using Beltran_Up_29 NE signature (8). **D,** Heatmap of single-sample GSVA pathway enrichment scores from bomedemstat-treated tumors compared to vehicle control using multiple NE signatures (8),(48),(63),(64),(65–67). **E,** *ASCL1* Log_2_ FPKM via bulk RNA-seq in bomedemstat-treated LTL331R tumors. **F,** qPCR for *ASCL1* mRNA expression in NCI-H660 treated *in vitro* with 10nM iadademstat. Relative mRNA expression changes calculated using 2^(-ΔΔCT)^ method with GAPDH as loading control. Significance assessed using two-tailed unpaired *t*-test. **G,** Representative IHC images of ASCL1 expression from end-of-study LTL331R PDXs treated with vehicle or bomedemstat *in vivo* and **H,** quantified using QuPath (53). Significance assessed using two-tailed unpaired *t*-test. Scale bars, 50µm. **I,** Western blot for ASCL1 protein expression in LTL331R spheroids treated with vehicle or bomedemstat [0.01-1.0µM] for 3 days *in vitro*. Representative blots from three independent experiments. **J,** ASCL1 after stable transduction of LTL331R with either empty vector (EV) control or ASCL1-OE plasmid treated with vehicle or 10nM iadademstat for 3 days *in vitro*. **K,** LTL331R EV control and ASCL1-OE proliferation measured after 14 days using CellTiter-Glo 3D. **L,** LTL331R EV control and ASCL1-OE treated with 100nM bomedemstat after 14 days normalized to vehicle control using CellTiter-Glo 3D. **M,** LTL331R ASCL1-OE cell proliferation after 10-100nM bomedemstat treatment over 12 days using CellTiter-Glo 3D. Cell proliferation reported as relative light units (RLU). For all assays, cells initially seeded at 10,000 cells/well. Significance assessed using two-tailed unpaired *t*-test. **N,** LTL331R ASCL1-OE sphere-formation images captured at day 12 after 10nM bomedemstat treatment. **O,** ASCL1 expression in NCI-H660 after transduced with ASCL1-OE plasmid or ASCL1 stable shRNA-mediated knockdown (ASCL1-KD) compared with EV control. **P,** Cell proliferation of NCI-H660 EV, ASCL1-OE, and ASCL1-KD measured using CellTiter-Glo 3D after 14 days. Significance assessed using two-tailed unpaired *t*-test.

To determine whether LSD1i-mediated suppression of ASCL1 in NEPC was required for growth suppression, we constitutively overexpressed *ASCL1* (ASCL1-OE) in LTL331R spheroids grown *ex vivo* or in NCI-H660 spheroids (Fig. 2J, O) and tested response to LSD1i. Although ASCL1-OE enhanced proliferation compared to empty-vector (EV) controls (Fig. 2K), cells remained sensitive to bomedemstat treatment (Fig. 2L). Treatment of ASCL1-OE cells with LSD1i suppressed ASCL1 protein abundance to levels at or below the vehicle-treated controls, and yet there was still no rescue in growth (Fig. 2J, L). Dose-dependent loss of cell proliferation and sphere-forming ability was also observed in ASCL1-OE cells treated with bomedemstat over 12 days (Fig. 2M, N). Although NEPC growth was augmented by ASCL1-OE, it was not impaired by *ASCL1* shRNA-mediated knockdown (Fig. 2O, P), suggesting that ASCL1 is dispensable for NEPC proliferation, and its downregulation is not required for LSD1i-mediated efficacy in these NEPC models. Taken together, these results indicate that the NE transcriptional profile is sustained by LSD1 and that small molecule inhibitors of this enzyme downregulate ASCL1 and the NE transcriptional profile, though this is not their sole mechanism of efficacy. LSD1-targeted therapy in NEPC may therefore trigger the activation of alternative non-NE transcriptional programs in the absence of ASCL1.

### LSD1 inhibition induces YAP1 expression in NEPC

Previous temporal motif analyses studying the ARPC-NEPC transdifferentiation trajectory have described transcriptional drivers from early to late-stage mCRPC tumors (6). Separate work has classified SCLC into subtypes based on transcription factors that are either NE drivers (ASCL1, NEUROD1) or non-NE drivers (YAP1, POU2F3) (7). Because LSD1i attenuated ASCL1 expression and the NE transcriptional program in these NEPC models, we explored the possibility that non-NE drivers defining different tumor transdifferentiation stages may be induced and activated by this therapy. We considered well-characterized transcription factors involved in adenocarcinoma-to-NE transdifferentiation (6,7,24,25). Among non-NE transcription factors, *YAP1* and *TEAD2* emerged as the top-enriched transcripts induced by LSD1i in LTL331R PDXs (Fig. 3A). *YAP1* was also among the top 30 globally upregulated genes in these treated PDXs (Fig. S4A). Expression of YAP1 is canonically silenced in PDXs from NEPC but retained in PDXs representing the three other mCRPC subtypes (Fig. 3B), consistent with prior reports (12). *YAP1* upregulation by LSD1i in NEPC models was confirmed by qPCR in LTL331R and NCI-H660 (Fig. 3C). Using the NCI-H660 ASCL1-KD model (Fig. 2O), loss of ASCL1 was sufficient to increase *YAP1* expression (Fig. 3D), suggesting that LSD1i-mediated suppression of ASCL1 also drives upregulation of *YAP1*. Furthermore, LSD1i led to elevated YAP1 protein abundance *in vitro* by bomedemstat or iadademstat in a dose-dependent manner (Fig. 3E, Fig. S4B) and *in* vivo, as shown by IHC (Fig. 3F, G). Downstream YAP1 transcriptional activity was also enriched in LSD1i-treated LTL331R PDXs, as assessed by GSEA (Fig. 3H, S4C) or qPCR for target genes including CYR61 (*CCN1*) (Fig. S4D). Several *TEAD* signatures were also significantly enriched *in vivo* following bomedemstat treatment in LTL331R tumors (Fig. 3I).

**Figure 3.**
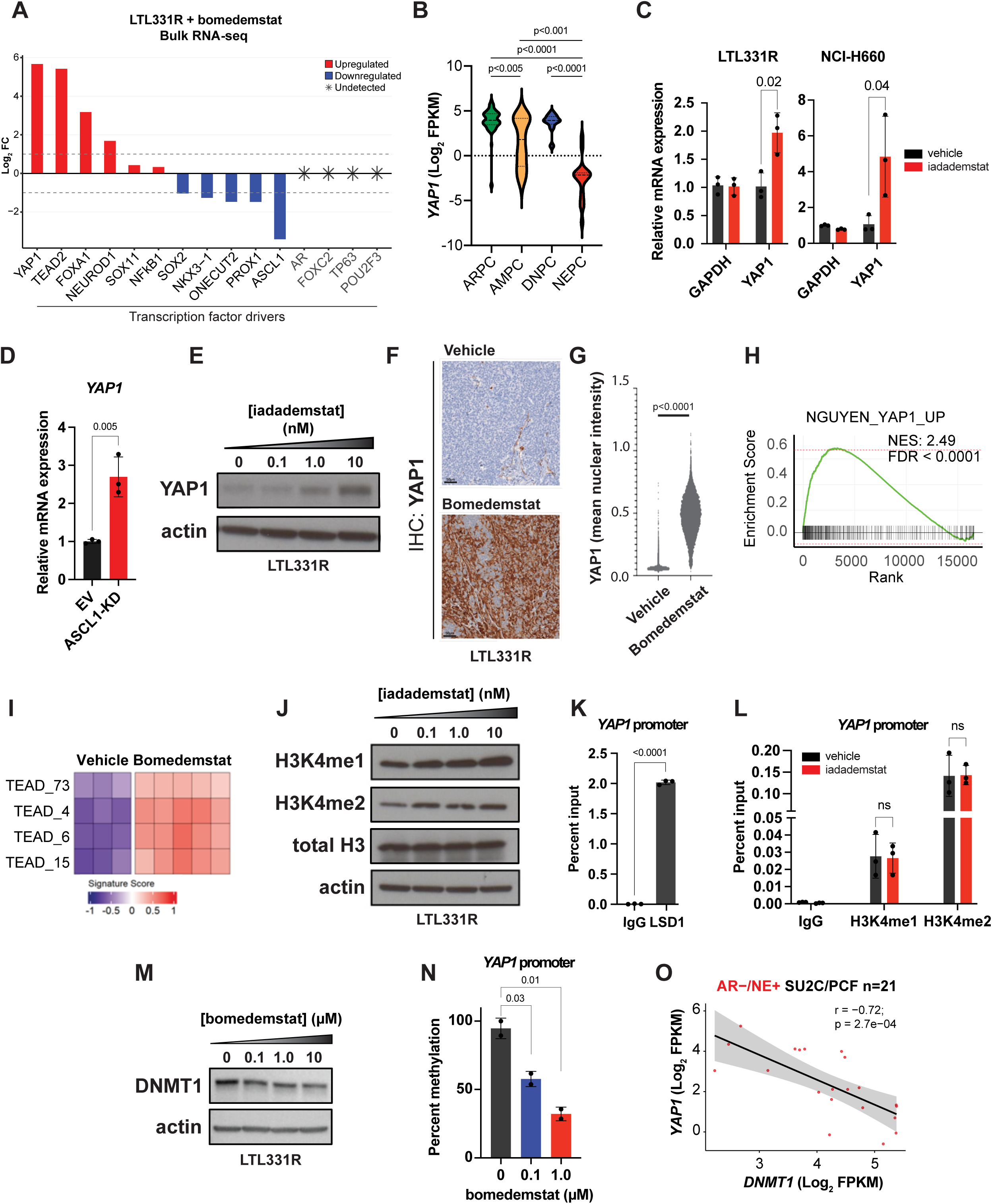
LSD1 inhibition induces YAP1 expression in NEPC. **A,** Transcript expression for transdifferentiation transcription factor drivers analyzed from RNA-seq on bomedemstat-treated LTL331R tumors. **B,** YAP1 expression across large PDX cohort (n = 135) of clinically relevant pathologic subtypes including ARPC (n = 93), AMPC (n =13), NEPC (n = 18), and DNPC (n =11). P-values determined by Kruskal-Wallis test followed by Tukey method. **C,** qPCR for *YAP1* mRNA expression in LTL331R and NCI-H660 treated *in vitro* with vehicle or 10nM iadademstat. **D,** qPCR for *YAP1* mRNA expression in the NCI-H660 ASCL1-KD model compared to control. Relative mRNA expression changes calculated using 2^(-ΔΔCT)^ method with *GAPDH* as loading control. Significance assessed using two-tailed unpaired *t*-test. **E,** Western blot for YAP1 protein abundance after vehicle or 0.1-10nM iadademstat treatment compared to vehicle. Representative blots from three independent experiments. **F,** Representative IHC images of YAP1 expression from end-of-study tumors from LTL331R PDXs treated with vehicle or bomedemstat *in vivo* and **G,** quantified using QuPath (53). Significance assessed using two-tailed unpaired *t*-test. Scale bars, 50µm. **H,** GSEA using Nguyen YAP1_UP (68) signature and **I,** TEAD activity signatures (Table S1) from bomedemstat-treated LTL331R tumors with GSVA scores. **J,** Western blot for H3K4me1 and H3K4me2 using total histone H3 and actin as loading controls after 3 days of treatment with vehicle or 0.1-10nM iadademstat. Representative blots from three independent experiments. **K,** ChIP LSD1 or matched IgG at *YAP1* promoter followed by qPCR in LTL331R spheroids. **L,** ChIP H3K4me1, H3K4me2, or matched IgG at *YAP1* promoter followed by qPCR in LTL331R spheroids treated with vehicle or 0.1-10nM iadademstat after 3 days. Calculated by percent input method. Significance assessed using two-tailed unpaired *t*-test. **M,** DNMT1 protein abundance following bomedemstat-treated LTL331R spheroids. **N,** Percent of *YAP1* promoter methylated in LTL331R spheroids following vehicle or 100-1000nM bomedemstat-treatment (n=2). Significance assessed using two-tailed unpaired *t*-test. **O,** *YAP1* vs *DNMT1* polyA FPKM transcript abundance in mCRPC patient data from SU2C/PCF dataset (13) using cBioPortal. Pearson r and p-value shown on plot.

To elucidate how LSD1i leads to de-repression of YAP1, we interrogated LSD1 recruitment to the *YAP1* promoter and the abundance of its canonical histone substrates. We hypothesized that LSD1 suppresses *YAP1* expression by demethylating the active histone marks, H3K4me1/2, and that LSD1i impedes this activity, leading to H3K4me1/2 accumulation at the *YAP1* promoter. We confirmed by Western blot that global H3K4me1/2 abundance was augmented in iadademstat-treated LTL331R spheroids (Fig. 3J). We then performed chromatin immunoprecipitation (ChIP)-qPCR on LSD1 and H3K4me1/2 at *YAP1*. We found that the *YAP1* promoter was enriched for both LSD1 (Fig. 3K) and its substrates H3K4me1/2 (Fig. 3L). However, despite LSD1 localization to the *YAP1* promoter, H3K4me1/2 abundance was not augmented by LSD1i (Fig. 3L). This suggested that LSD1 may regulate YAP1 expression through a noncanonical mechanism in NEPC. Previous work has demonstrated that LSD1 regulates the stability of non-histone proteins *via* post-translational modifications (26). We also considered a prior report demonstrating that *YAP1* silencing is mediated by DNA methylation (12). Importantly, DNA methyltransferase 1 (DNMT1) is one non-histone substrate stabilized by LSD1 (26,27). Indeed, LTL331R spheroids treated with LSD1i led to a dose-dependent reduction in DNMT1 protein (Fig. 3M) but not mRNA transcript abundance (Fig. S5A). Next, we asked whether LSD1i affected DNA methylation at the *YAP1* promoter. Here, we observed that LSD1i significantly decreased *YAP1* promoter DNA methylation levels with increasing inhibitor concentrations (Fig. 3N). To assess whether global DNA methylation levels were altered by LSD1i as a secondary effect, we performed liquid chromatography using mass spectrometric (LC-MS/MS) detection using genomic DNA from LTL331R spheroids treated with bomedemstat or decitabine as a positive control. Global DNA methylation levels were reduced with decitabine treatment, as expected, but unaltered by bomedemstat (Fig. S5B), suggesting locus-specific regulation by LSD1. Furthermore, in analyzing mCRPC tumors from the SU2C/PCF dataset, we determined that high expression of *DNMTs* (*DNMT*-*1*, *-3A*, *-3B*) is associated with low expression of *YAP1* exclusively in the NEPC patient samples (Fig. 3O, S6A-D). These *YAP1*-low/*DNMT1*-high tumors also presented with elevated expression of *KDM1A* as compared to the other mCRPC subtypes from this clinical dataset (Fig. S7A). Taken together, these data suggest that LSD1i alleviates YAP1 repression in NEPC by modulating DNA methylation through DNMT1 in a locus-specific manner.

### YAP/TEAD disruption enhances the antitumor efficacy of LSD1-targeted therapy

YAP1 has been shown to exert oncogenic or tumor-suppressor functions depending on the cancer type. In prostate and lung adenocarcinomas YAP1 is expressed and oncogenic, whereas in small cell neuroendocrine carcinomas YAP1 is silenced and tumor-suppressive (28). These classes, which are termed “YAP^ON^” or “YAP^OFF^”, have been extended to various types of solid tumors based on their YAP1 functional status (28). We confirmed that in the absence of LSD1i, YAP1 is silenced and that genetic overexpression of YAP1 (YAP1-OE) in LTL331R leads to growth suppression (Fig. 4A, B), thereby confirming that this NEPC model behaves as a YAP1^OFF^ tumor type.

**Figure 4.**
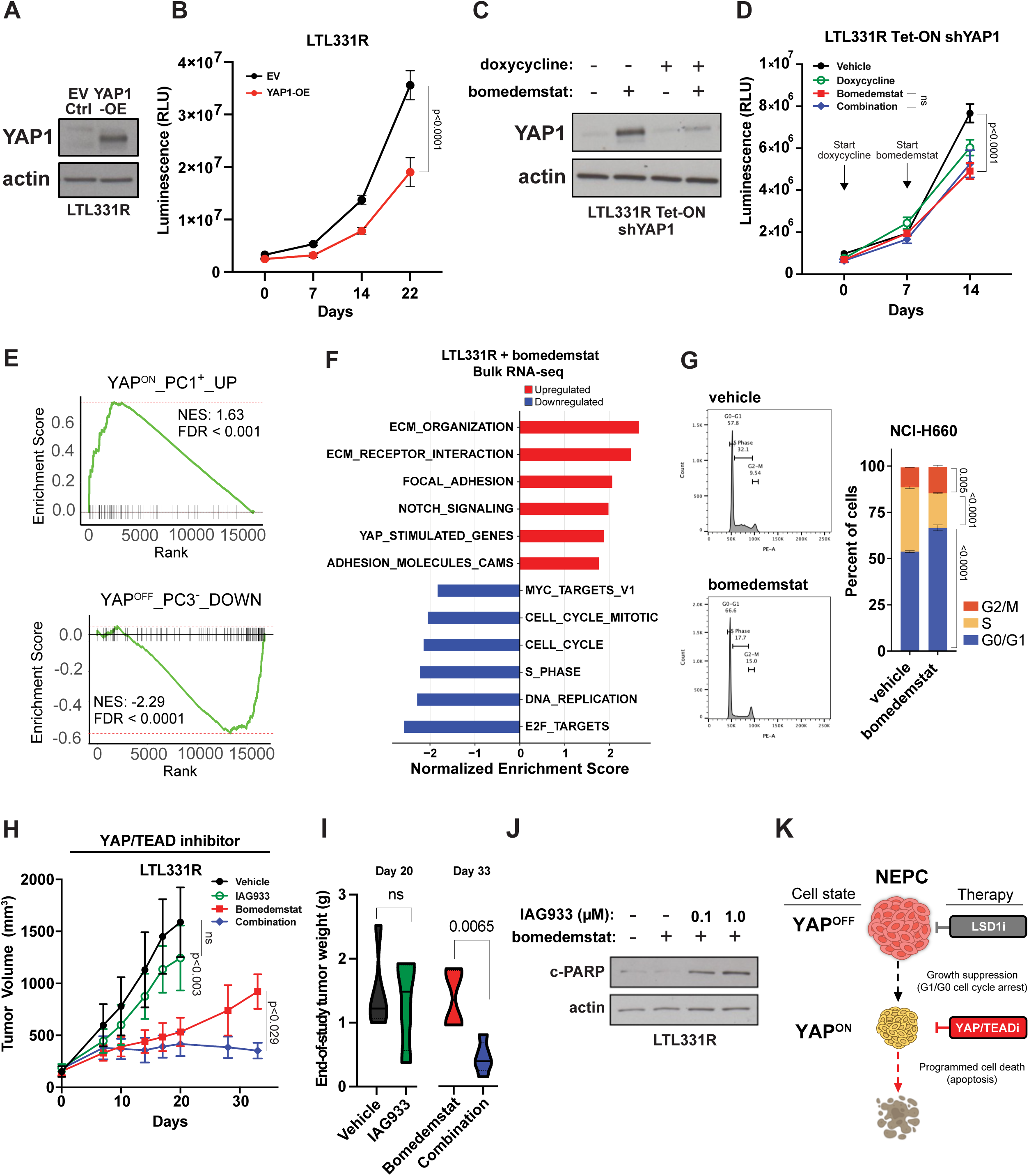
LSD1 inhibitor-treated NEPC is vulnerable to YAP/TEAD disruption. **A,** Western blot for YAP1 expression after stable transduction of LTL331R with YAP1-OE plasmid. **B,** YAP1-OE LTL331R cell proliferation compared to EV control after 22 days using CellTiter-Glo 3D from 10,000 cells seeded. Significance assessed using two-tailed unpaired *t*-test. **C,** Western blot for LTL331R transduced with doxycycline-inducible Tet-ON system targeting *YAP1* via shRNA-mediated knockdown (LTL331R Tet-ON shYAP1). Successfully transduced cells were pre-treated with 1µg/mL doxycycline for 7 days prior to addition of 100nM bomedemstat or vehicle to assess knockdown efficiency. **D,** Cell proliferation of LTL331R Tet-ON shYAP1 treated with 100nM bomedemstat after 7 day 1µg/mL doxycycline pre-treatment assessed using CellTiter-Glo 3D from 5,000 cells seeded. Significance assessed using two-tailed unpaired *t*-test. **E,** GSEA for YAP1^ON^ cancer class signature (PC1^+^_UP) (28) in LTL331R PDXs after bomedemstat treatment *in vivo*. GSEA for YAP1^OFF^ cancer class signature (PC3^-^_DOWN) (28) in LTL331R PDXs after bomedemstat treatment *in vivo*. **F,** Normalized enrichment scores for YAP1^ON^-associated pathways up- or down-regulated by bomedemstat in NEPC tumors. Selected HALLMARK_, KEGG_, or REACTOME_ pathways with an adjusted p-value<0.01 and FDR<0.1 are shown. **G,** Flow cytometric cell cycle analysis using propidium iodide fluorescence-activated cell sorting (FACS) of NCI-H660 spheroids treated with 100nM bomedemstat after 25 days, n=3 independent experiments. Significant differences between vehicle-treated or bomedemstat-treated cells in G0/G1-, S-, or G2/M-phase determined using two-way ANOVA with Šidák correction for multiple comparisons. **H,** Male NSG mice bearing LTL331R tumors response to bomedemstat monotherapy (LSD1i, dosed PO daily at 40 mg/kg), IAG933 monotherapy (YAP/TEADi, dosed PO daily at 240mg/kg), bomedemstat/IAG933 combination therapy with respective doses, or vehicle control (n=4-6/group). Data presented as the mean ±SEM; p-values determined using linear mixed model (LMM) with Šidák correction for multiple comparisons. **I,** Final tumor masses (g) measured for each group at end-of-study (Day 20 and Day 33). For vehicle-treated tumors, n=5; IAG933-treated tumors, n=4; bomedemstat-treated tumors, n=4; combination-treated tumors, n=5. Significance assessed using two-tailed unpaired *t*-test. **J,** Western blot for cleaved PARP abundance in LTL331R after 7 day treatment with bomedemstat +/- 0.1 or 1μM IAG933. **K,** Schematic illustration depicting LSD1i therapy causing growth arrest of NEPC and conversion of this subtype from a YAP^OFF^ to a YAP^ON^ cell state with enhanced sensitivity to YAP/TEAD-targeted therapy.

Given that YAP1 exhibits a tumor suppressive function in NEPC and that inhibiting LSD1 induces its expression, we initially hypothesized that post-translational stabilization of YAP1 would enhance growth suppression in NEPC tumors in combination with LSD1i. Strategies to stabilize tumor suppressor proteins are being explored clinically for several cancer types (29,30). Serine-threonine kinases 3/4 (STK3/4) are Hippo-signaling effector proteins whose activation of LATS1/2 leads to phosphorylation of YAP1, which is then sequestered into the cytoplasm and subsequently degraded (31). Inhibition of STK3/4 therefore reduces YAP1 degradation, permitting its translocation into the nucleus for downstream YAP1 transcriptional activity. Using SBP-3264, a small molecule inhibitor of STK3/4 (32), we treated NSG mice bearing LTL331R PDXs to determine antitumor response when combined with LSD1i. As expected, SBP-3264 alone exerted no efficacy due to absence of LSD1i-mediated upregulation of YAP1 expression (Fig. S8A, B). Surprisingly, however, LTL331R PDXs remained insensitive to SBP-3264 even when combined with bomedemstat or iadademstat (Fig. S8A, B). Instead, the role of YAP1 appeared context-dependent. Although NEPC growth was impaired by ectopic *YAP1* expression (Fig. 4B), these results indicated that the antitumor efficacy by LSD1i occurred independently of endogenous *YAP1* induction. To genetically test this hypothesis, we transduced LTL331R spheroids with a Tet-ON doxycycline-inducible shRNA targeting *YAP1* and subsequently treated this model with bomedemstat to determine effects on growth (Fig. 4C). Here, the antiproliferative effect induced by bomedemstat in NEPC was not abrogated by knockdown of *YAP1*, confirming that LSD1i-mediated growth suppression is independent of endogenous *YAP1* induction despite its activity as a tumor suppressor in this cancer type (Fig. 4D). Therefore, although YAP1 expression and activity is induced by LSD1i in NEPC, the growth suppressive ability of YAP1 is not retained in this context.

This suggested an intriguing possibility in which LSD1 regulates not only YAP1 expression, but also function in NEPC tumors. Although YAP1 acts as a tumor suppressor in NE tumors, LSD1i may reprogram NEPC from a YAP^OFF^ to a YAP^ON^ state, thereby converting YAP1 into a targetable dependency. Therefore, we examined our bulk RNA-seq data from LTL331R PDXs treated with bomedemstat to determine whether tumors were enriched for signatures defining these YAP^ON^/ YAP^OFF^ states (28). Here, we observed enrichment of the YAP^ON^ signature and downregulation of the YAP^OFF^ signature in LSD1i-treated NEPC tumors (Fig. 4E). Other YAP1-regulated processes including ECM, focal adhesion, and NOTCH signaling pathways were also enriched in these tumors as anticipated in YAP^ON^ cancers (Fig. 4F). Importantly, YAP1 gene pathways that drive tumor growth were not activated in LSD1i-treated tumors. Oncogenic YAP^ON^ pathways such as DNA replication, cell cycle transition, E2F, and MYC activity were dramatically reduced following LSD1i (Fig. 4F). These RNA-seq data were supported using flow cytometric cell cycle analysis which demonstrated that NEPC cells were arrested in G0/G1 phase after 25 days of bomedemstat treatment (Fig. 4G). Rather than acting as an oncogenic driver of aggressive disease, as is the case for conventional YAP^ON^ cancer types, YAP1 pathway activation appears to sustain survival of NEPC tumors under LSD1i-treatment pressure.

Given this treatment-induced YAP^ON^ state, we hypothesized that targeting YAP1 would only exert antitumor efficacy in NEPC when combined with LSD1i. IAG933 is a clinically-investigated small molecule inhibitor of YAP1 activity that disrupts interaction with its protein partners TEAD1-4, which commence transcriptional effector function with YAP1 (33). NSG mice bearing the LTL331R model treated with IAG933 alone showed no significant difference between vehicle-treated groups as expected (Fig. 4H). Strikingly, however, we observed significant growth suppression in LTL331R tumors when IAG933 was administered in combination with bomedemstat compared to either agent as a monotherapy (Fig. 4H). Tumor mass was significantly reduced in the combination group compared to LSD1i or YAP/TEAD disruption alone (Fig. 4I). Cell death was also activated using this combination therapy as indicated by the elevated abundance of the apoptosis marker, cleaved PARP (Fig. 4J). Furthermore, dual inhibition of LSD1 and YAP1 was generally well-tolerated in mice with dose de-escalation over the duration of study (Fig. S9A), indicating both efficacy and safety considerations of this treatment strategy are achieved. Altogether, these data demonstrate LSD1i epigenetically switches NEPC from a YAP^OFF^ to a YAP^ON^ state (Fig. 4K), thereby sensitizing these treatment-induced tumors to YAP/TEAD disruption.

## Discussion

In this study, we determined that 8 out of 12 mCRPC CDX/PDX models were responsive to LSD1i using bomedemstat. A prior report identified enhanced sensitivity to LSD1i in mCRPC models with *RB1* loss (20). Our data classifying C4-2B with functional *RB1* as insensitive to LSD1i is consistent with their results. We also extend their findings by identifying *TP53* LOF as a potential biomarker of response across different mCRPC models representing the four major subtypes – ARPC, AMPC, DNPC, and NEPC. In our study, *RB1* status alone would have excluded several mCRPC models responsive to LSD1i including VCaP, LAPC4, CWR22-RH, and PC3 despite their observed sensitivity to bomedemstat *in vivo*. PC3 sensitivity to LSD1i *in vivo* is also consistent with a prior report using this model (21). Meanwhile, mCRPC models with *TP53* LOF showed consistent sensitivity to LSD1i. This effect is illustrated in differential response to LSD1i in CWR22-RH and 22Rv1 which are derived from the same parental model, CWR22. LSD1i sensitivity is observed in CWR22-RH, which exhibits *TP53* G154F LOF whereas non-responsive 22Rv1 expresses functional *TP53*. Our data demonstrate a strong correlation between LSD1i response across 12 mCRPC models and *TP53* status. Consistent with our findings, gastric cancer cell models harboring *TP53* LOF are growth impaired by the reversible LSD1i GSK690, whereas those expressing wildtype *TP53* are intrinsically resistant to this therapy (34). In prostate cancer, LSD1 expression is inversely correlated with *TP53* signaling in multiple patient cohorts (9), further supporting genomic stratification methods for LSD1i. Yet, whether *TP53* LOF is required for sensitivity to LSD1i in mCRPC and if induction of wildtype *TP53* reverses this effect should be the subject of investigation for future work. Nevertheless, *TP53* LOF/deficiency frequently co-occurs with *RB1* loss in a majority of NEPC tumors suggesting LSD1i therapy has the potential to be especially effective for these patients with very poor clinical outcomes and a lack of robust treatment options. This is substantiated by the exceptional antitumor response to LSD1i observed in the NEPC models.

Several clinical trials are currently exploring LSD1-targeted monotherapies in a variety of diseases including two unreported trials enrolling prostate cancer patients. NCT04628988 was designed to investigate the effects of pulrodemstat in resensitizing mCRPC patients to anti-hormone therapy. A dual LSD1/HDAC6 inhibitor is being studied in NCT05268666 for patients with advanced solid tumors which included NEPC in the eligibility criteria for part 2 of the study focused on NE-like/NE-derived tumors. However, not all LSD1 inhibitors share the same clinical potential. For instance, bomedemstat is well-tolerated by patients in multiple Phase 2 clinical trials including for AML (NCT02842827) and is currently in Phase 3 testing for essential thrombocythemia (NCT06079879). Iadademstat also demonstrates an overall tolerable drug profile as reported from Phase 1 trial results (35) and is entering Phase 1/2 for SCLC (NCT06287775) and AML (NCT06357182). Conversely, GSK2879552, has not been pursued further clinically due to limited efficacy and major toxicities of thrombocytopenia and encephalopathy (NCT02034123) (36). Seclidemstat (SP2577) is being investigated for myelodysplastic syndrome (MDS) and chronic myelomonocytic leukemia (CMML) (37) (NCT04734990) despite the FDA placing a partial hold on the trial due to unexpected adverse effects; the trial has since resumed.

As these agents advance in the clinic, a deeper understanding into how LSD1i affects tumors remains important to identify patients most likely to benefit, mitigate adverse effects, and enhance efficacy. Our transcriptional characterization of NEPC in response to LSD1i revealed marked suppression of ASCL1 and significant downregulation of the NE transcriptional profile. Other studies using SCLC models demonstrate iadademstat-mediated LSD1i leads to loss of NE gene signatures and suppression of ASCL1 (22,23). Despite this, our data indicated that ASCL1 downregulation appears dispensable for LSD1i-mediated suppression of NEPC proliferation as assessed by LSD1i treatment of LTL331R ASCL1-OE. Likewise, knockdown of ASCL1 did not abrogate proliferation in NCI-H660. Although ASCL1 is important for initiating the NE transcriptional program as others have shown (38), our data is consistent with reports demonstrating that established NEPC tumors will continue to grow even in the absence of ASCL1 (24). Independence from ASCL1 may occur due to a variety of factors that promote NEPC survival. In the context of LSD1i, our data suggest that NEPC undergoes epigenetic reprogramming to a transcriptional class defined by activation of YAP1 whose expression is canonically silenced in this mCRPC subtype (12). Among transcription factors that define the adenocarcinoma-to-NE lineage trajectory, YAP1 and TEAD2 were notably enriched by LSD1i *in vivo*, suggesting that YAP1 induction is preferred by NEPC when ASCL1 is absent.

Furthermore, we demonstrate that LSD1i-mediated regulation of YAP1 occurs via DNA methylation by DNMT1. Despite LSD1 occupancy at the *YAP1* promoter, direct histone demethylation of H3K4me1/2 at the locus is unchanged by LSD1i. Instead, DNA methylation is reduced with bomedemstat treatment, indicating LSD1 post-translationally stabilizes DNMT1. Our data suggest stability of DNMT1 is diminished by bomedemstat, leading to its degradation and subsequent locus-specific hypomethylation. Although it is unclear how global DNA hypomethylation is circumvented when LSD1 is inhibited, compensatory mechanisms may occur through other DNMTs, such as DNMT3A/B, which are involved in *de novo* hypermethylation (39–41). Additionally, epigenetic homeostasis may be maintained by feedback loops that reduce the expression or activity of Ten-eleven translocation (TET) enzymes, which demethylate DNA (42). This mechanism may also be preserved in SCLC which is defined by distinct ASCL1 or non-NE YAP1 subclasses and shares molecular features with NEPC (43). How LSD1 regulates ASCL1 is yet to be determined. In SCLC, LSD1i-mediated suppression of ASCL1 is observed with activation of NOTCH signaling (22). Our data indicate that NOTCH signaling is enriched in bomedemstat-treated LTL331R tumors (Fig. 4F), suggesting its upregulation may downregulate ASCL1 expression, assuming this mechanism is conserved across NE carcinomas. However, further work is necessary to clarify ASCL1 regulation by LSD1 in the context of NEPC.

YAP1 function is context-dependent, acting as an oncogene in prostate adenocarcinoma but exhibiting a tumor suppressive role in NEPC (28). This functional dichotomy suggested that LSD1i-mediated YAP1 activation was either responsible for suppressing NEPC growth or, in the absence of ASCL1, promoting cancer cell survival. Evidence for the latter was substantiated with *in vivo* data demonstrating that dual inhibition of LSD1 and YAP/TEAD pathway activation enhanced antitumor efficacy greater than either monotherapy. In this study, we utilized IAG933 which inhibits YAP pathway activation by disrupting the protein-protein interaction between YAP1 and TEAD transcription factors (33). This inhibitory mechanism may be essential for enhanced antitumor efficacy given our Tet-ON inducible YAP1-KD data demonstrating no difference in NEPC growth when LSD1i was added to doxycycline-treated cells (Fig. 4D). Since TEADs are able to recognize and bind YAP1 or its homolog TAZ to drive YAP pathway activation (44), our data suggests that disabling promiscuous TEAD activity may be more effective than inhibiting YAP1 alone in the context of LSD1i.

Consistent with the role of epigenetic regulation in defining YAP pan-cancer classes (28), LSD1 appears critical for defining this state in NEPC. Modulating the YAP^ON^/YAP^OFF^ cancer classes by targeting LSD1 has implications beyond prostate cancer. Sensitization to YAP1-targeted therapies using LSD1i may also be effective in other YAP^OFF^ solid tumors such as SCLC, neuroblastoma, low-grade gliomas, retinoblastomas, NE colorectal, and NE pancreatic cancers. Further studies are warranted to understand this functional switch in YAP^ON^/YAP^OFF^ tumors. Taken together, we report LSD1i as an effective therapy in *TP53* LOF NEPC models which activate a non-NE YAP^ON^ transcriptional profile vulnerable to YAP/TEAD inhibition, thereby potentially improving outcomes for patients diagnosed with this lethal subtype.

## Materials and Methods

### Cell lines and reagents

All cell lines were obtained from ATCC unless otherwise noted. LNCaP, C4-2B, 22Rv1, DU145, and PC3 were grown in RPMI-1640 (Thermo Fisher, Waltham, MA) + 10% FBS (Corning Cellgro, Corning, NY). LAPC4 were grown in Iscove’s Modified Dulbecco’s Medium (IMDM, Thermo Fisher) + 5% FBS supplemented with 1 nM R1881 (Perkin-Elmer, Boston, MA). LN-95 is an androgen-independent cell line derived from LNCaP as previously reported (45) and cultured in phenol red-free RPMI-1640 supplemented with 10% charcoal-stripped FBS (Hyclone Laboratories Inc, Logan, UT). NCI-H660(46,47) were grown in advanced DMEM/F12 (Gibco, Gaithersburg, MD) supplemented with B27 (Gibco), EGF (10 ng/mL, Sigma), and FGF (10 ng/mL, Sigma). The LTL331 PDX(48) was generously provided by Dr. Yuzhou Wang, which we converted to spheroid culture grown in advanced DMEM/F12 (Gibco) supplemented with B27 (Gibco), EGF (10 ng/mL, Sigma), +/- FGF (10 ng/mL, Sigma). HEK293-T cells were grown in Dulbecco’s Modified Eagles Medium (DMEM) + 10% FBS. All cultures were also supplemented with 1% L-glutamine and 1% Penicillin-Streptomycin. All cultures were maintained under standard tissue culture conditions in a humidified incubator at 37°C with 5% CO2 and were passaged using 0.05% Trypsin-EDTA twice weekly for adherent models or biweekly for spheroid models. All lines were confirmed to be mycoplasma negative, routinely tested using the MycoSensor PCR Assay kit (Agilent Technologies, La Jolla, CA) and genetically authenticated within the last year via STR profiling performed in the Johns Hopkins Genetic Resource Core Facility.

### Plasmids/transductions

HEK293-T cells were seeded in a T150 flask in complete growth medium (DMEM + 10% FBS). 24 hrs later, medium was replaced with opti-MEM (Addgene). Viral particles were prepared by combining 15 µg of the indicated plasmid, 30 µg psPAX2 (Addgene), 7.5 µg pMDG (Addgene), 132 µL FuGENE HD Transfection Reagent (Promega), and 2.25 mL opti-MEM and gently mixing. The mixture was added to cells and incubated at 37°C for 72hrs. Viral particles in the supernatant were collected and spun down for 10 min at 1,000 rpm. Supernatant was filter-sterilized through a 0.45 µM filter. PEG solution (51 mL 50% PEG 6000 [Sigma-Aldrich], 21.7 mL 4 M NaCl, 23.3 mL PBS) was added to the supernatant at a 1:2 ratio and incubated at 4°C for 24 hrs. Mixture was centrifuged for 45 min at 5000 rpm at 4°C, and the precipitate was resuspended in 1 mL HBSS. Viral titer was checked according to the qPCR Lentivirus Titer Kit (abm, Richmond, Canada). For transduction of target cells, viral particles were added to T25 flasks at 10 MOI in the presence of 8 µg/mL polybrene, and media was exchanged after 24 hrs. After another 24 hrs, puromycin (5 µg/mL) was added for selection. LTL331R and NCI-models grown as spheroids were transduced with viral particles as bulk populations. <u>Plasmids</u>: *EF.CMV.RFP* (empty vector control; Addgene #10879), *EF.CMV.RFP_ASCL1* (full length hASCL1 cloned into EcoRV site of EF.CMV.RFP vector), *EF.CMV.RFP_YAP1* (full length hYAP1 cloned into EcoRV site of EF.CMV.RFP vector), *pLKO.1_EV* (empty vector control; Addgene #10879), *pLKO.1_shASCL1* (shRNA targeting hASCL1 cloned into pLKO.1_EV), *pLKO-Tet-On-YAP1-shRNA1* (Addgene #176846). *EF.CMV.RFP* was a gift from Linzhao Cheng (49) (Addgene plasmid #17619; http://n2t.net/addgene:17619 ; RRID:Addgene_17619). *pLKO.1-TRC* control was a gift from David Root (50) (Addgene plasmid #10879; http://n2t.net/addgene:10879 ; RRID:Addgene_10879). *pLKO-Tet-On-YAP1-shRNA1* was a gift from Roland Friedel (51) (Addgene plasmid #176846; http://n2t.net/addgene:176846;RRID:Addgene_176846).

### Western blot

Cell pellets were collected and lysed using a buffer composed of NP-40 (Thermo Fisher Scientific), cOmplete protease inhibitor (Roche), and PhosSTOP phosphatase inhibitor (Roche). Reaction was incubated at 4°C for 15 min and sonicated. Soluble fraction was collected before normalization based on protein concentration (15-50µg) using Pierce™ BCA Protein Assay (Cat. #23227) and mixed with 2x Laemmli buffer (50 mM Tris-HCl, pH 6.8, 2% SDS, 10% Glycerol, 0.005% Bromophenol blue). Samples were denatured at 95°C for 4 min before SDS-PAGE. Denatured samples were separated using 4–15% Criterion™ TGX™ Precast Protein Gel and transferred to PVDF membrane. Membranes were blocked in 5% milk (TBS + 0.1% Tween-20) for 1 hour and incubated with primary antibody shaking at 4°C overnight followed by respective secondary antibody the following day shaking for 1 hour at room temperature. ECL™ or SuperSignal™ West Femto Maximum Sensitivity Substrate was used for development by Cytiva Amersham Hyperfilm™ films which were digitally scanned. <u>Antibodies:</u> Anti-LSD1 (Santa Cruz #sc-53875), anti-ASCL1 (BD Biosciences, #556604; Santa Cruz #sc-374104), anti-YAP1 (CST #14047), anti-DNMT1 (Sigma #MABE306, clone DNM-2C1), anti-H3K4me1 (CST #5326), anti-H3K4me2 (EpiCypher #13-0027), anti-total histone H3 (CST #9715), anti-cleaved PARP (CST #5625), anti-β-actin (Sigma-Aldrich #A5441), anti-mouse HRP secondary (CST #7076), anti-rabbit HRP secondary (CST #7074), or anti-rat HRP secondary (CST #7077).

### Chromatin-immunoprecipitation (ChIP)

1x10^6 cells were collected for each ChIP reaction and processed according to the ab500 ChIP Kit (Abcam). Briefly, cells were fixed using 1.1% formaldehyde, quenched with glycine, and lysed using manufacturer’s buffers. Chromatin was sonicated and run on 1% agarose gel to confirm DNA fragment size of 200-500 bp. Samples were incubated at 4°C overnight with ChIP-grade antibodies for LSD1 (Abcam #ab17721; 4µg), H3K4me1 (CST #5326; 1:50), H3K4me2 (CST #9725; 1:50), in addition to IgG control (4µg). The next day antibody bound-DNA was purified and quantified by qRT-PCR using percent input method. Primers used for *YAP1* promoter – FP: GGGAGTGTGCAGGAATGTAGCAAC, RP: CGACTGAGACAGAAACTCGCC.

### qRT-PCR

mRNA expression was determined by quantitative reverse transcription (qRT-PCR) according to standard methods. Briefly, RNA was isolated from a known quantity of cells using the Qiagen RNeasy kit. cDNA was generated using iScript cDNA synthesis kit (Bio-Rad). PCR amplification of cDNA performed using iTaq Universal SYBR Green Supermix (Bio-Rad) according to manufacturer’s instructions with the following IDT pre-designed PrimeTime™ Primers: *ASCL1* – Hs.PT.56a.23464.gs, *YAP1* – Hs.PT.58.14881945, *DNMT1* – Hs.PT.58.28037916, *CCN1* – Hs.PT.58.27029250.gs, *HPRT1* – Hs.PT.58v.45621572, *GADPH* – Hs.PT.39a22214836.

### Cell proliferation and sphere formation

NCI-H660 and LTL331R (with indicated vectors) were seeded at 2,500 or 10,000 cells/well, respectively, in a 96-well plate format. Unless otherwise stated, treatment with either bomedemstat or iadademstat began on Day 0 along with initial measurement in proliferation. Proliferation was measured using CellTiter-Glo® 3D Cell Viability Assay (#G9681) according to manufacturer’s recommendation with luminescence measured as relative light units (RLU). Celigo Imaging Cytometer was used to assess spheroid formation with images taken on Day 12 of cell treatment.

### Xenograft and patient-derived xenograft (PDX) models

All animal procedures were approved by the Johns Hopkins University School of Medicine Institutional Animal Care and Use Committee. Adult (>8 wk old) male NOD-*scid* IL2Rgamma^null^ (NSG) mice were obtained from the Sidney Kimmel Comprehensive Cancer Center (SKCCC) Animal Core Facility and used for xenografting. Adult male NSG mice were inoculated subcutaneously (1-2x10^6^) in 200 µL 50% Matrigel (n=3-8/grp). Male NSG mice were castrated contingent on type of CDX/PDX inoculated. The following models were xenografted into castrate males: C4-2B, LN-95, CWR-22RH, 22Rv1, LvCaP-2R, PC3, and LTL331R. The following models were xenografted into intact males: LNCaP, VCaP, LAPC4, DU145 and NCI-H660. Castration was performed according to standard protocols by making a scrotal incision under anesthesia.

When tumors became palpable (∼100-200mm^3^), PDX-bearing mice were dosed with: bomedemstat (LSD1i from Imago Biosciences, a subsidiary of Merck) at 40 mg/kg/day PO, iadademstat (LSD1i from Oryzon Genomics) at 0.4 mg/kg/week PO, SBP-3264 (STK3/4i from Selleckchem) at 10 mg/kg for 5 days ON and 2 days OFF per week IP, IAG933 (YAP/TEADi from Novartis) at 240mg/kg/day PO. Dosing schedule and method was identical in vehicle controls (bomedemstat – 200 µL 0.5% carboxymethylcellulose, 0.5% Tween 80, 5% DMSO; iadademstat – 200 µL saline; SBP-3264 – 10% Tween-80, 5% DMSO, 85% dH2O; IAG933 – 0.5% methylcellulose + 0.1% Tween80 in 100mM Phosphate Buffer pH 8. For combination groups, SBP-3264 was started after 7 day lead-in with bomedemstat or iadademstat; IAG933 was started after 1 day lead-in with bomedemstat. Following lead-in, dosing schedule and method were consistent when used in combination studies. Tumor volume (length x width x height x *0.5326*) was measured every 3-5 days via calipers as previously described (52). Body weights monitored longitudinally as indicated. Tumor weight was recorded at the end-of-study when indicated.

### Histology and immunohistochemistry (IHC)

H&E performed by the Oncology Tissue Services Core of Johns Hopkins University according to validated standard protocols. For immunohistochemical staining, slides were deparaffinized and steamed for 30 min in 10 mmol/L sodium citrate (pH = 6, Vector Labs). Primary antibodies and dilutions used were as follows: ASCL1 (Abcam ab211327, 1:50) and YAP1 (Abcam ab52771, 1:50). Immunocomplexes were detected using anti-Rabbit IgG (Leica Microsystems Cat. PV6119) with DAB as the chromogen. Tissue sections were counterstained with hematoxylin and slides were digitized on a Ventana DP 200 Slide Scanner (Roche). Image analyses were performed using QuPath (53).

### Flow cytometric cell cycle analysis

After trypsinization/neutralization of NCI-H660 spheroids, isolated single cells were counted and normalized to 500,000 cells per condition. Cells were centrifuged at 500xg for 5 minutes and washed with HBSS. Washed cells were then fixed dropwise with pre-chilled 70% ethanol and stored at -20°C until analysis. On the day of flow cytometric analysis, the ethanol was removed after centrifugation at 500xg, and the cells were washed twice with PBS. They were then stained with a solution containing 50 µg/mL propidium iodide (Sigma-Aldrich) and 100 µg/mL RNase A (Sigma-Aldrich). The cells were acquired on a BD FACSCelesta flow cytometer and analyzed using FlowJo software version 10.4.2.

### RNA sequencing (RNA-seq) analysis

Total RNA from flash frozen PDX tumors was extracted and DNase-treated using the RNeasy Mini kit (Qiagen). RNA concentration, purity, and integrity was assessed by NanoDrop (Thermo Fisher Scientific) and Agilent TapeStation. RNA-seq libraries were constructed from 1µg total RNA using the Illumina TruSeq Stranded mRNA LT Sample Prep Kit according to the manufacturer’s protocol. Barcoded libraries were pooled and sequenced by Illumina NovaSeq 6000 generating 50 bp paired end reads. Sequencing reads were mapped to the hg38 human genome and mm10 mouse genomes using STAR.v2.7.3a (54). All subsequent analyses were performed in R. Sequences aligning to the mouse genome deriving from potential contamination with mouse tissue were removed from the analysis using XenofilteR (55) (v1.6). Gene level abundance was quantitated using the GenomicAlignments (56). Differential expression between groups was assessed using limma (57), filtered for a minimum expression level using the filterByExpr function with default parameters prior to testing, and using the Benjamin-Hochberg false discovery rate (FDR) adjustment. Genome-wide gene expression results were ranked by their limma statistics and used to conduct Gene Set Enrichment Analysis (GSEA) (58) to determine patterns of pathway activity utilizing the curated pathways from within the MSigDBv7.4. Single sample enrichment scores were calculated using GSVA (59) with default parameters using genome-wide log2 FPKM values as input. GSVA scores of groups displayed in boxplots were compared using two-sided Wilcoxon rank tests.

### Site-specific DNA methylation analysis

A previously validated assay combining methylated-DNA precipitation and methylation-sensitive restriction enzyme digestion (COMPARE-MS) was used for site-specific DNA methylation analyses (60). In brief, DNA samples were digested with AluI and HhaI (New England Biolabs) and methylated DNA fragments were enrichment using recombinant MBD2-MBD (Clontech) immobilized on magnetic Tylon beads (Clontech). The precipitated DNA containing methylated DNA fragments was eluted and subjected to quantitative real-time PCR using IQ SYBR Green Supermix (Bio-Rad) with primers specific to the second intron of *YAP1* (hg38 chr11:102,112,102-102,112,272) – FP: 5’-CGCTTTACAGTACCGAGGTTT; RP: 5’-GCCATGTTCATATCCGCAATAAG. For quantitative assessment of locus-specific methylation levels, CT-values of the samples of interest were normalized to CT-values of the positive control (*in vitro* fully methylated male genomic DNA) and methylation indices were calculated (ranging from 0-100%).

### Global DNA methylation mass spectrometry

LTL331R spheroids were treated with vehicle, 1µM bomedemstat, or 10µM decitabine for 7 days before DNA was extracted. DNA samples were digested and analyzed by LC-MS/MS for quantitation of 2dC, 5Me2dC, 2dG, and 5hm2dC based on a previously described method with the following changes (61). The LC system consisted of a Shimadzu UPLC system flowing into a Waters Nova Pak Silica column (3.9 × 150 mm, 4 µm) maintained at 40°C. Mobile phase A consisted of 5 mM ammonium acetate in water, and mobile phase B consisted of 5 mM ammonium acetate in water/acetonitrile (2:98, v/v). Gradient elution was performed at a flow rate of 1.4 mL/min with a total run time of 10 minutes using the following gradient conditions: 97% B at 0–3.2 min, 90% B at 7.2 min, 2% B at 7.3 min, and re-equilibration to 97% B at 8.0 min.

Mass spectrometric detection was carried out using an AB Sciex 6500+ triple quadrupole mass spectrometer (Framingham, MA, USA) with SelexION technology operating in positive electrospray ionization and multiple reaction monitoring (MRM) mode. The settings of the mass spectrometer for all analytes were as follows: curtain gas 10, collision gas 12, IonSpray voltage 5500 V, probe temperature 350 °C, GS1 40, GS2 30, and EP of 10 V. The optimized compound-dependent parameter values were as follows: DP values were 160 V for 2dC, 86 V for 5Me2dC, 176 V for 2dG, 100 V for 5hm2dC. CXP values were 4 V for 2dC, 8 V for 5Me2dC, 10 V for 2dG, 12 V for 5hm2Dc. CE values were 13 V for 2dC, 29 V for 5Me2dC, 15 V for 2dG, 15 V for 5hm2Dc.

The MRM *m/z* transitions monitored were: m/z 228.1→112.0 for 2dC, 242.1→126.1 for 5Me2dC, 268.5→152.1 for 2dG, and 257.8→141.8 for 5hm2dC, along with their corresponding isotope-labeled internal standards. The dwell time was set at 100 msec for all transitions. Data acquisition and quantitative analysis were performed using Analyst software version 1.7.1 or higher with quadratic regression and 1/x² weighting.

Calibration standards and quality control samples were prepared in blank digest matrix containing deferoxamine mesylate, ammonium acetate, Trizma base, EDTA, and ZnCl_2_. Calibration ranges were 0.1–400 ng/mL for 5hm2dC, 0.25–1000 ng/mL for 5Me2dC, and 2.5–10,000 ng/mL for 2dC and 2dG. QC samples were prepared at low, medium, and high concentrations.

Samples were prepared using acetonitrile extraction containing stable isotope-labeled internal standards ([^13^C^15^N_2_]-2dC, [D_3_]-5Me2dC, [^13^C_10_,^15^N_5_]-2dG, and [D_3_]-5hm2dC).

### Clinical co-expression analysis

Transcript correlations from 208 metastatic prostate cancer patients with mRNA expression Z-scores relative to all samples (log FPKM capture) were analyzed from the SU2C/PCF (PNAS 2019) (13) dataset using cBioPortal (62).

### Statistical analysis

All statistical analyses were performed in R or using GraphPad Prism (v.6). Normally distributed data was subjected to a two-tailed unpaired Student’s *t*-test. Statistical significance for multigroup comparisons was assessed using two-way ANOVA (or Linear Mixed Model; LMM) followed by Šidák correction for multiple comparisons. Variables with repeated measurements were analyzed using LMM to take into account the possible within-subject correlation in the data. Error bars for all data denote mean ± SD or SEM as indicated in figure legends.

### Data and materials availability

The RNA-seq data for this publication has been deposited in NCBI’s Gene Expression Omnibus (GEO) and are accessible through accession number GSE234523 for the raw and processed mouse-sequence subtracted PDX data. Clinical trial expression data obtained from cBioPortal. All other data generated in this study are available upon request from the corresponding author.

## Author contributions

<u>Conception and design</u>: SMJ, HYRJr, JTI, WNB. <u>Development of methodology</u>: SMJ, RK, IMC, SLD, SK1, LA, DMR, JHB. <u>Acquisition of data</u>: SMJ, RK, SK1, IMC, SLD, TEGK, LA, DMR, BH, ZA, JHB. <u>Provided resources</u>: TL, IMC, PSN, ZA, JHB. <u>Analysis and interpretation of data</u>: SMJ, RK, IMC, BH, RAP, LAS, MCH, PSN, HYRJr, JTI, WNB. <u>Writing, review, and/or revision of the manuscript</u>: SMJ, RK, IMC, SK2, PSN, SRD, MCH, JHB, HYRJr, LAS, WNB. <u>Study supervision</u>: JTI, LAS, WNB. <u>Final approval</u> <u>of manuscript</u>: all authors.

## Acknowledgements

The authors thank Kristen Uchtmann and Dalton McLean at the William Ricke laboratory (UW-Madison) for training in subrenal capsule inoculations used to initiate the LTL331/R model. The authors would also like to acknowledge the following sources of financial support: Genentech Drug Metabolism and Pharmacokinetics Fellowship (SMJ), W81XWH2210118 (RK), HT94252310029 (RK), PhRMA Foundation Pre-Doctoral Fellowship Grant Number 131427 (TEGK), T32CA009071, (AM), V Foundation (AM), P50 CA097186 (PSN), R01 CA266452 (PSN), PC230420 (PSN), P01 CA298991 (PSN), R01 CA279993 (PSN), R50CA274336 (IMC), Department of Defense Award HT9425-23-1-0105 (SK2), R01 5R01CA243184 (SK2), Department of Defense CDMRP Award No W81XWH-17-1-0528 (WNB), NIH R01 CA255259 (WNB), NIH R01 CA279993 (WNB), V Foundation (WNB), monetary gift from Imago Biosciences, a subsidiary of Merck & Co, Inc. (WNB, JTI), and the NIH-SKCCC CCSG award P30 CA006973, which provides support for the SKCCC Oncology Tissue Services, Analytical Pharmacology Core, Animal Resources, Cell Imaging, and Genetic Resources Core Facilities.

The authors would also like to acknowledge the use of Claude Sonnet 4.5 (Anthropic) to assist in generating R scripts for data visualization. In specific, predetermined GSEA data for pathways were used as input to generate bar plots in R for interpretation of LSD1i-induced gene changes. Additionally, a heatmap was generated by Claude Sonnet 4.5 after we independently identified gene alterations across mCRPC models using multiple publicly available datasets and reports.

## Supplemental figure legends

**Figure S1.**
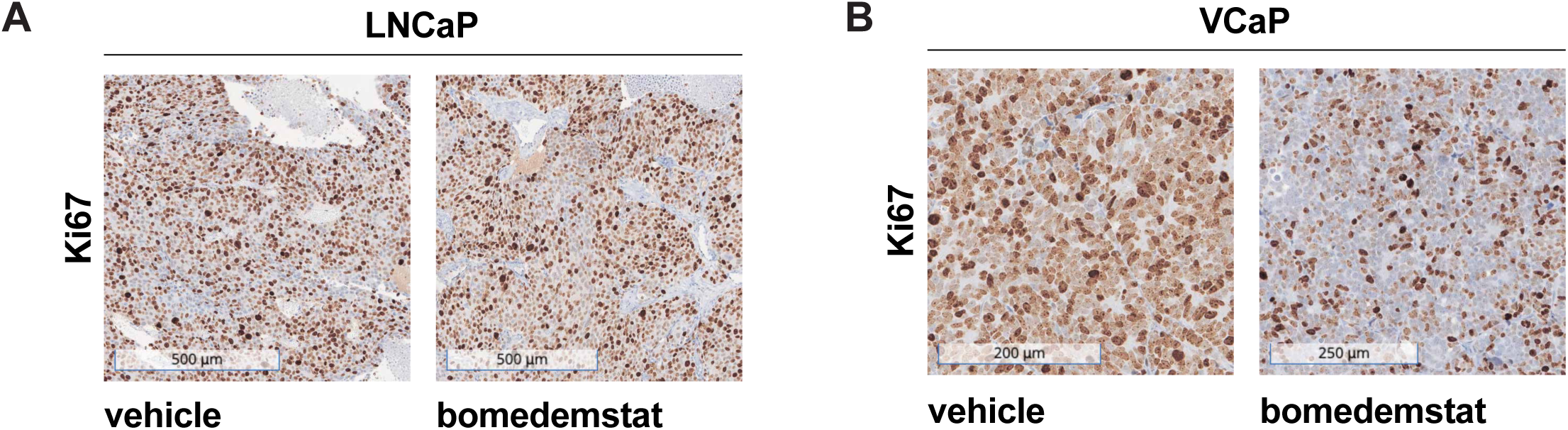
Immunohistochemistry for Ki67 in LSD1i-responder and non-responder models. **A,** Vehicle- or bomedemstat-treated end-of-study tumors processed for IHC and stained with Ki67 from LSD1i non-responder, LNCaP, or **B,** LSD1i-responder, VCaP.

**Figure S2.**
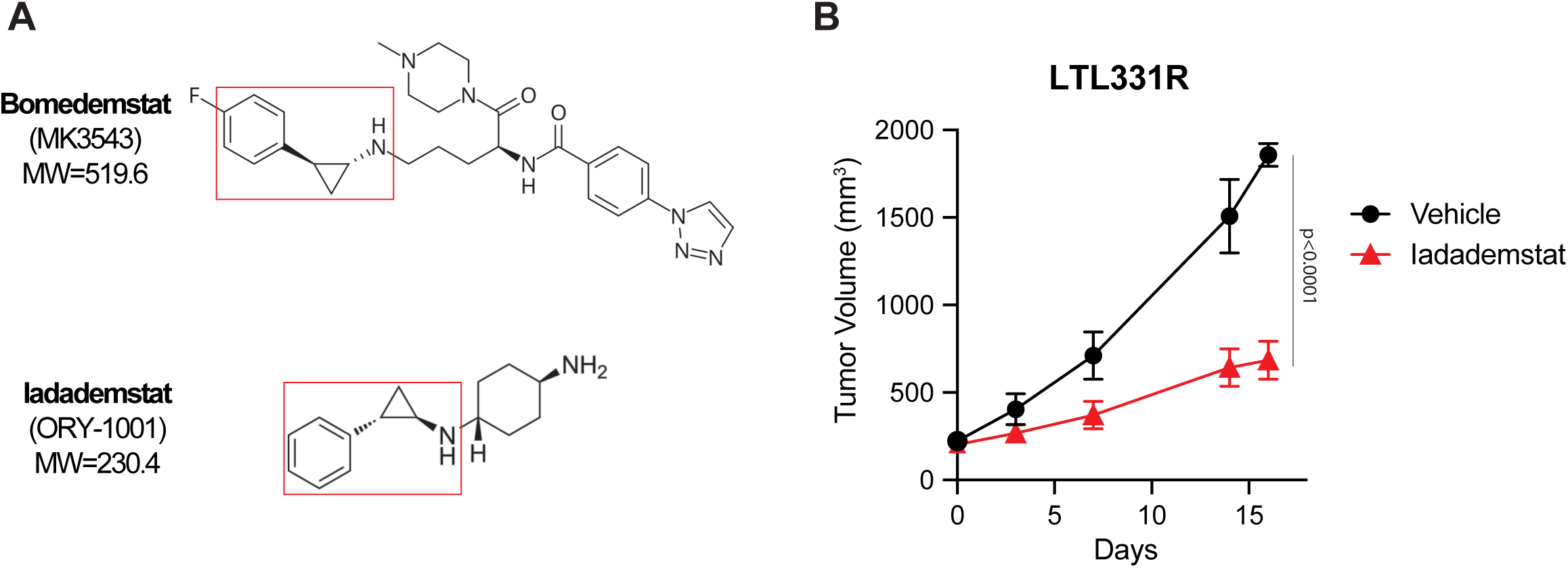
Antitumor efficacy by iadademstat in NEPC. **A,** Chemical structures for bomedemstat and iadademstat depicting conserved tranylcypromine group (outlined in red) that permits similar mechanism of action for LSD1i. **B,** Tumor growth curve for iadademstat dosed at 0.4 mg/kg/week PO compared to vehicle control in male NSG mice bearing LTL331R PDXs, n=5/group; data presented as the mean ± SEM and p-value determined using linear mixed model (LMM) with Šidák correction for multiple comparisons.

**Figure S3.**
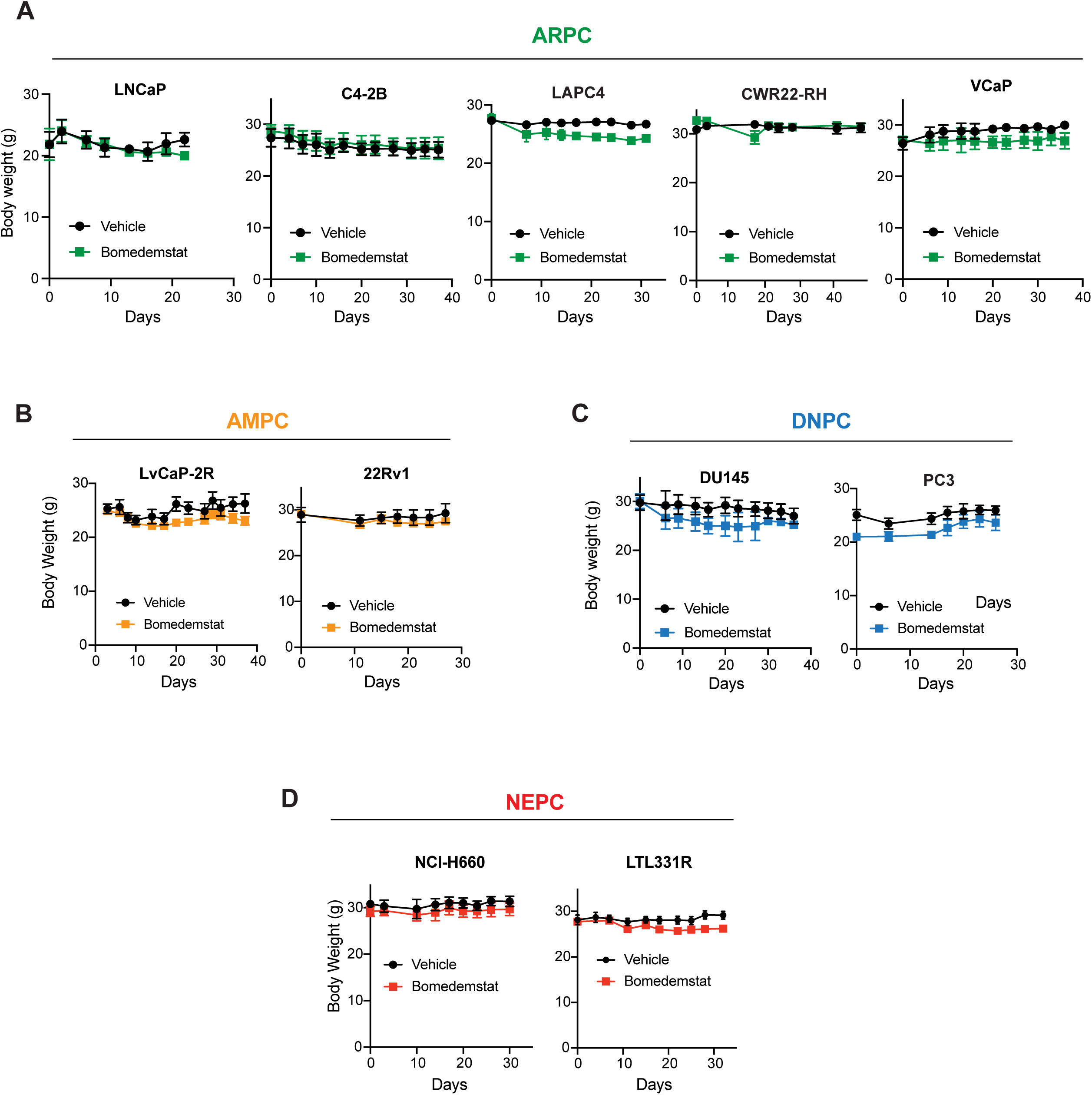
Body weights of mice bearing CDX/PDXs. **A,** Body weights recorded for mice dosed with bomedemstat (dosed PO daily at 40 mg/kg) or vehicle control in male NSG mice bearing models of ARPC: LNCaP (n=5), C4-2B (n=3-4), LAPC4 (n=4-5), CWR22-RH (n=3-4), VCaP (n=5), **B,** AMPC: 22Rv1 (n=5), LvCaP-2R (n=5-6), **C,** DNPC: PC3 (n=5), DU145 (n=5), and **D**, NEPC: NCI-H660 (n=5), LTL331R (n=5). Data presented as the mean ±SEM, n=mice per condition. Days on treatment shown.

**Figure S4.**
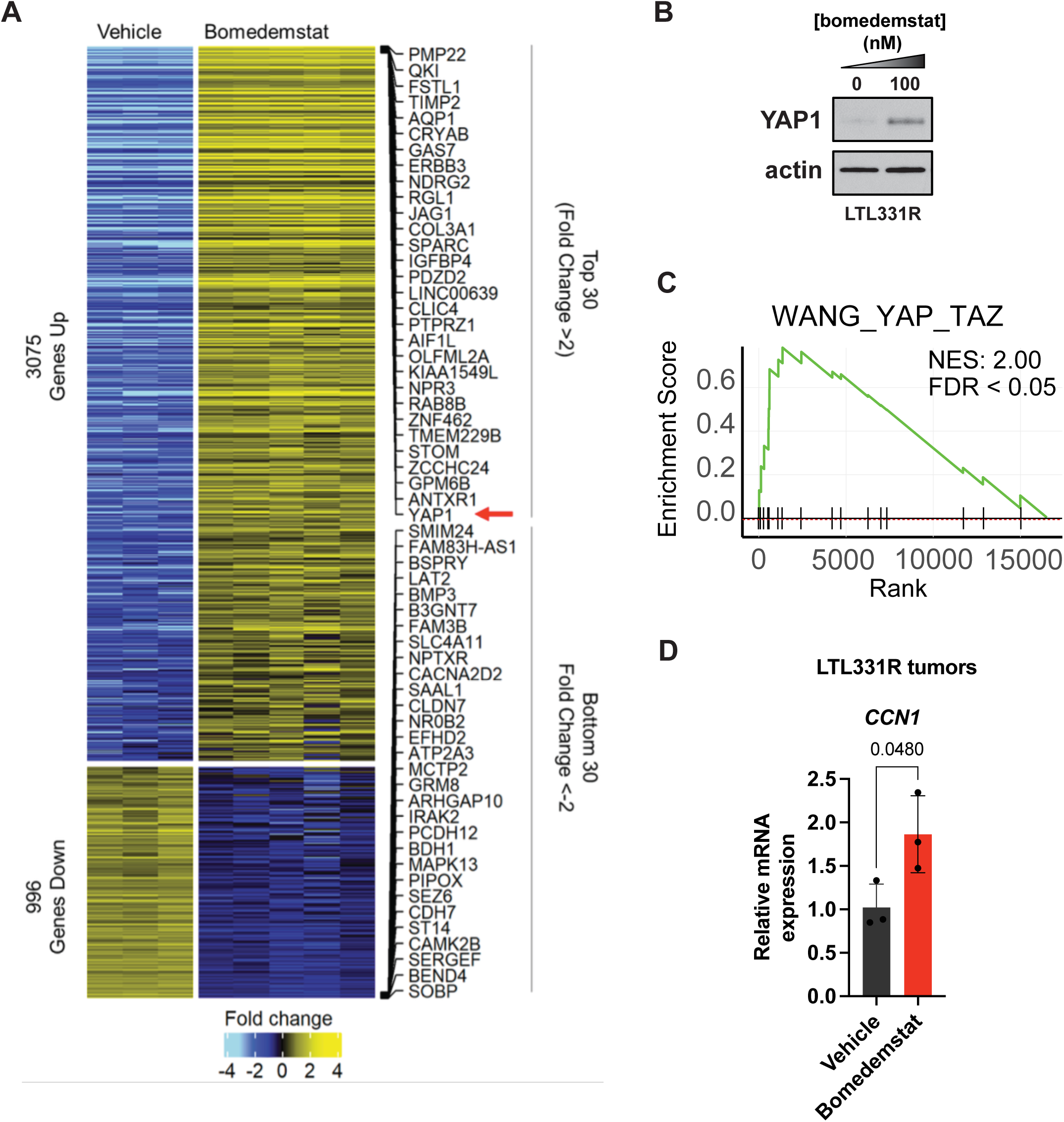
Transcriptional upregulation of YAP1 and its downstream targets by LSD1i. **A,** Top and Bottom 30 up- and down-regulated genes in LTL331R PDXs treated *in vivo* with vehicle (n=3) or bomedemstat (n=5). **B,** Western blot for YAP1 protein expression in LTL331R spheroids treated for 3 days with 100nM bomedemstat. **C,** YAP1 activity in LSD1i-treated LTL331R PDXs assessed by GSEA using Wang_YAP_TAZ signature (69). **D,** qPCR on *CYR61* mRNA expression in LTL331R PDXs treated with bomedemstat *in vivo*. Relative mRNA expression changes calculated using 2^(-ΔΔCT)^ method with *HPRT1* as loading control. Significance assessed using two-tailed unpaired *t*-test.

**Figure S5.**
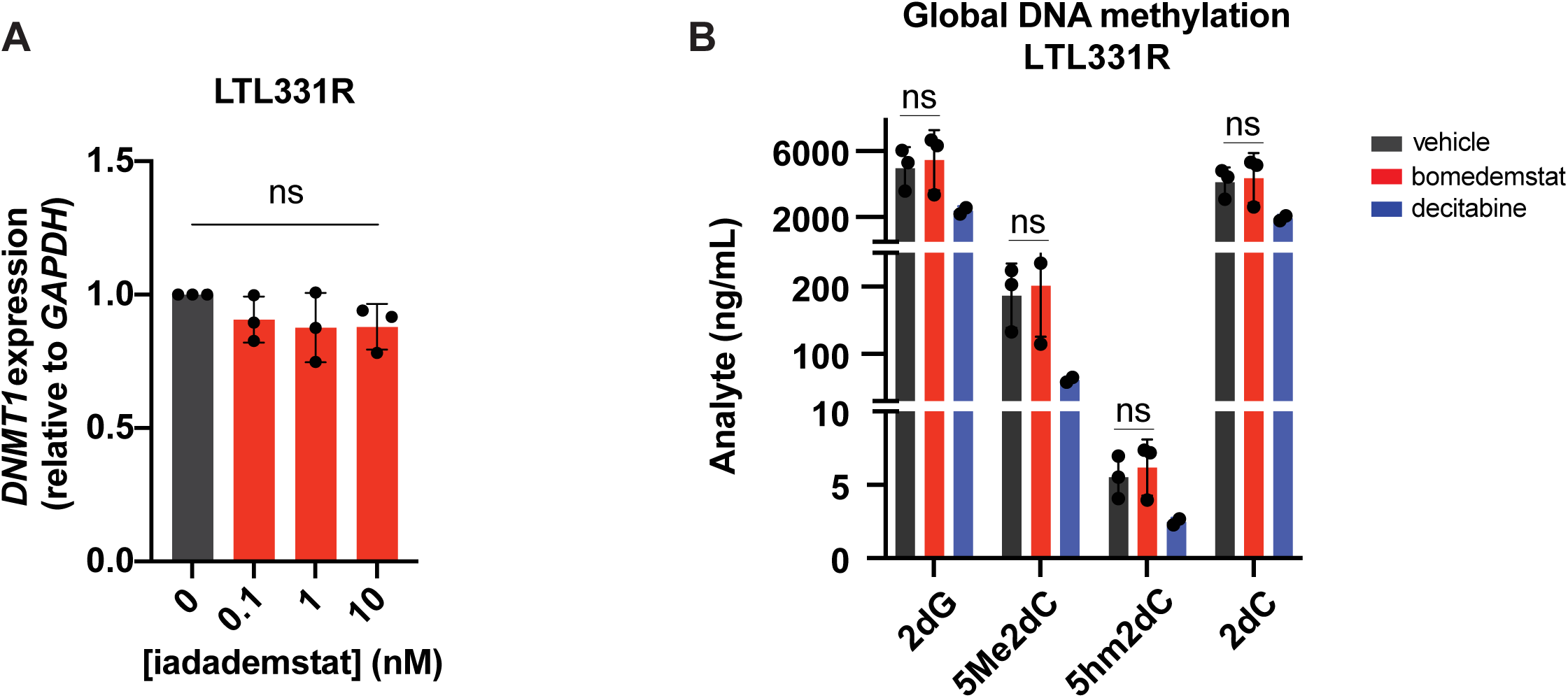
LSD1i-mediated effects on DNMT1 and DNA methylation. **A,** qPCR on DNMT1 mRNA expression in LTL331R spheroids after 3 days on 0.1-10nM iadademstat treatment. Relative mRNA expression changes calculated using 2^(-ΔΔCT)^ method with *HPRT1* as loading control. Significance assessed using two-tailed unpaired *t*-test. **B,** Global DNA methylation levels in LTL331R spheroids treated with 1µM bomedemstat or 10µM decitabine. Assessed by 5me2dC and 5hm2dC abundance via LC-MS/MS. Significance assessed using two-tailed unpaired *t*-test.

**Figure S6.**
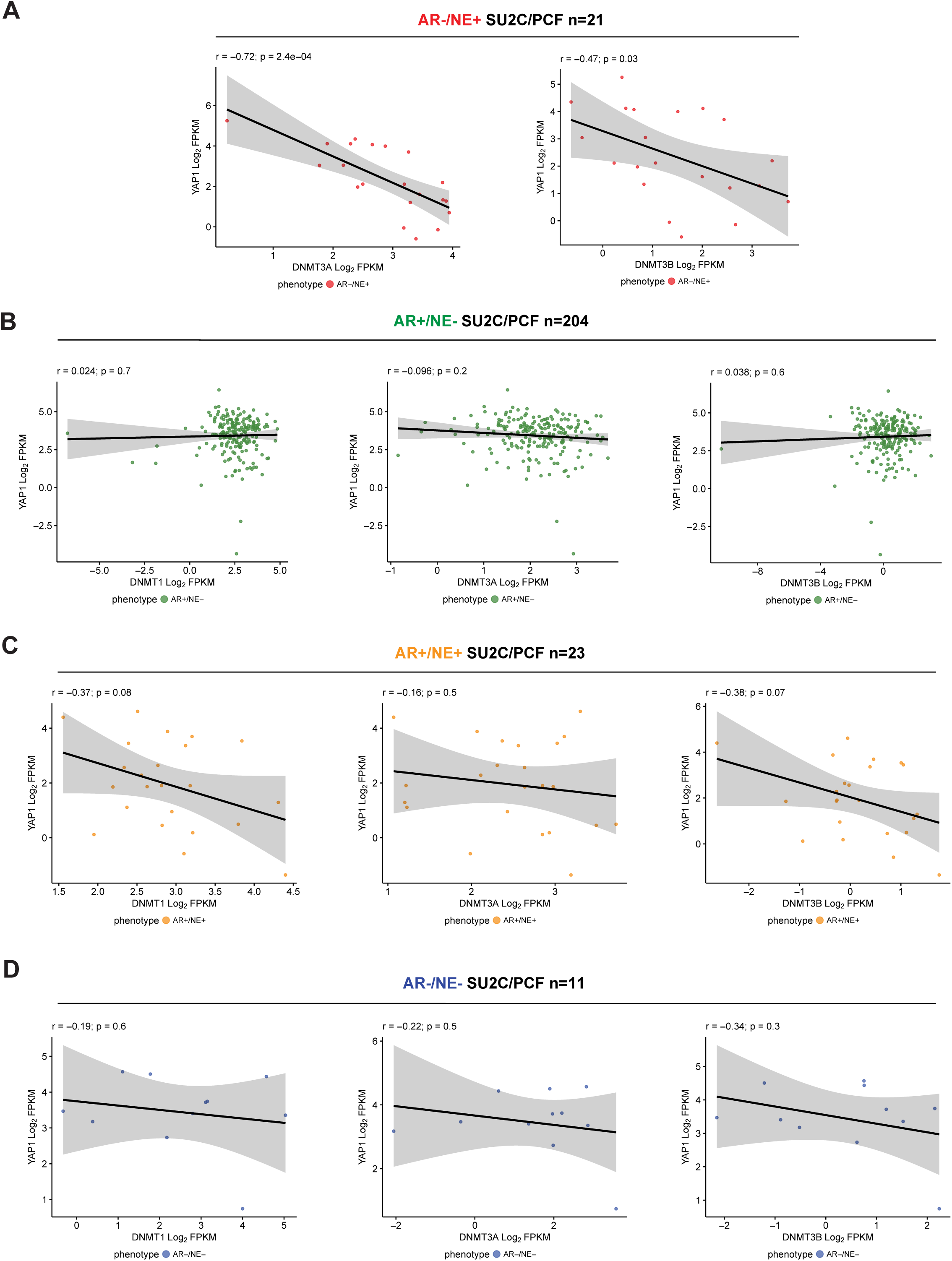
Clinical association between expression of *YAP1* and DNMTs in mCRPC subtypes. **A,** *YAP1* vs *DNMT-1, 3A,* or *3B* transcript abundance in NEPC, **B,** ARPC, **C,** AMPC, and **D,** DNPC patient samples from SU2C/PCF dataset (13). PolyA FPKM displayed for datasets with Pearson r and p-values shown on plots.

**Figure S7.**
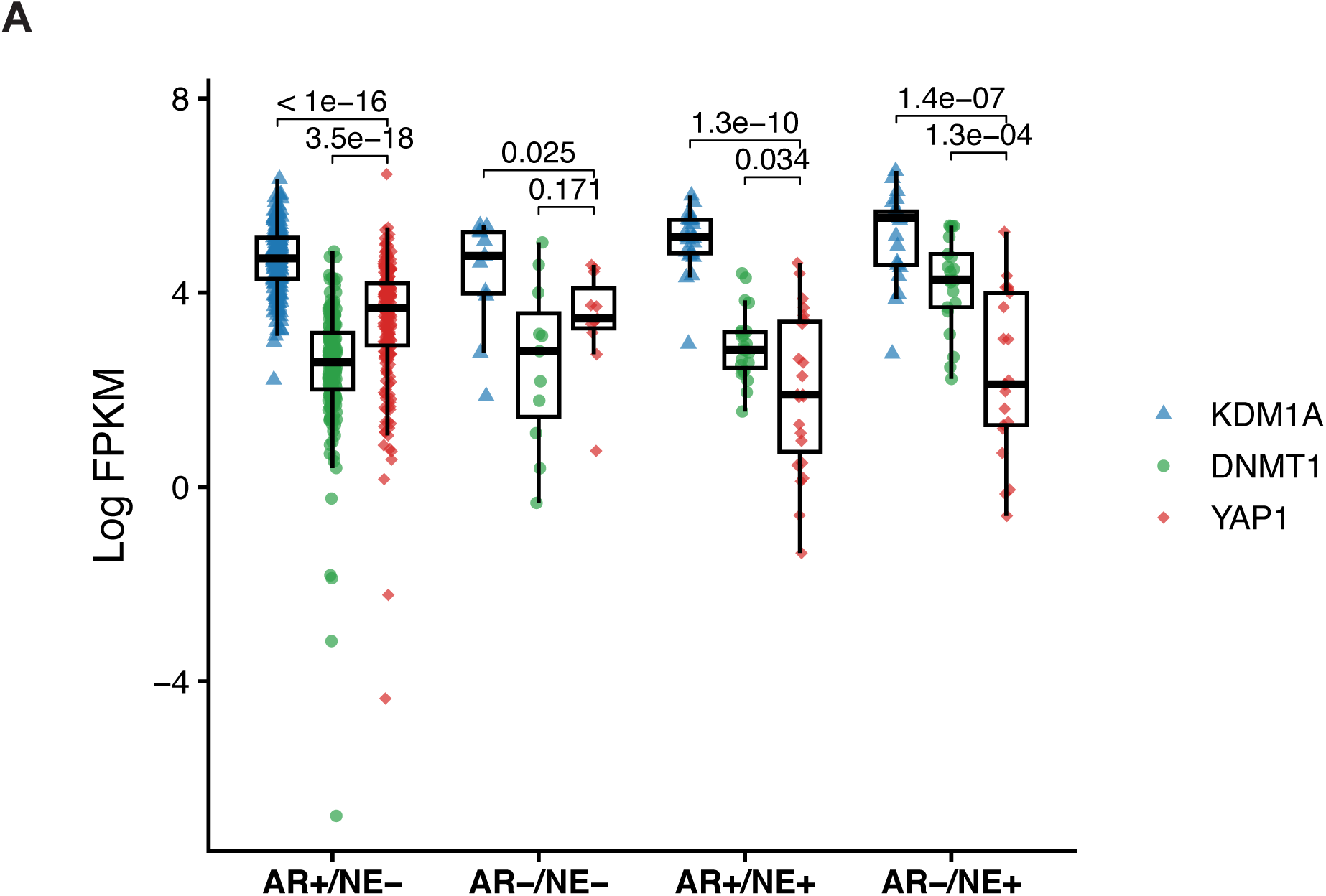
*KDM1A*, *DNMT1*, and *YAP1* correlation across mCRPC patient subtypes. **A,** *KDM1A, DNMT1*, and *YAP1* transcript abundance in patients samples from SU2C/PCF dataset (n=259) (13) stratified by AR+/NE-, AR-/NE-, AR+/NE+, or AR-/NE+ subtypes. PolyA FPKM displayed and groups compared by Wilcoxon-rank tests with pairwise BH adjusted P values (within subtype) shown on plots.

**Figure S8.**
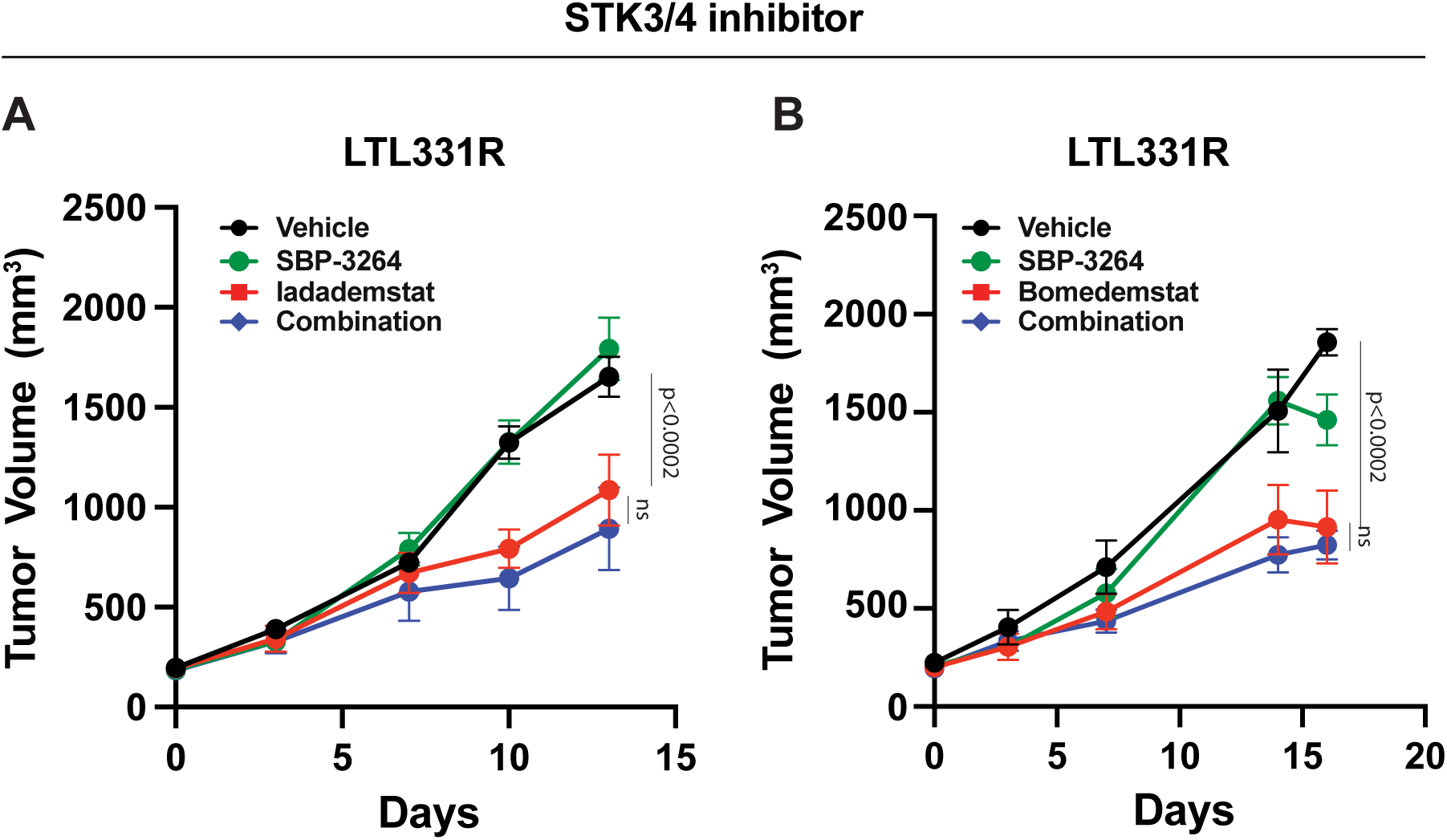
Dual inhibition of LSD1 and upstream YAP1 modulators STK3/4 in NEPC. **A,** Mice bearing LTL331R PDXs treated with vehicle, iadademstat (400µg/kg per week PO), SBP-3264 (10 mg/kg for 5 days ON and 2 days OFF per week IP), or iadademstat/SBP-3264 combination therapy with respective doses. **B,** Mice bearing LTL331R PDXs treated with vehicle, iadademstat (40mg/kg per day PO), SBP-3264 (10 mg/kg for 5 days ON and 2 days OFF per week IP), or bomedemstat/SBP-3264 combination therapy with respective doses. N=5 mice per group. Data presented as the mean ±SEM; p-value determined using ANOVA or LMM with Šidák correction for multiple comparisons.

**Figure S9.**
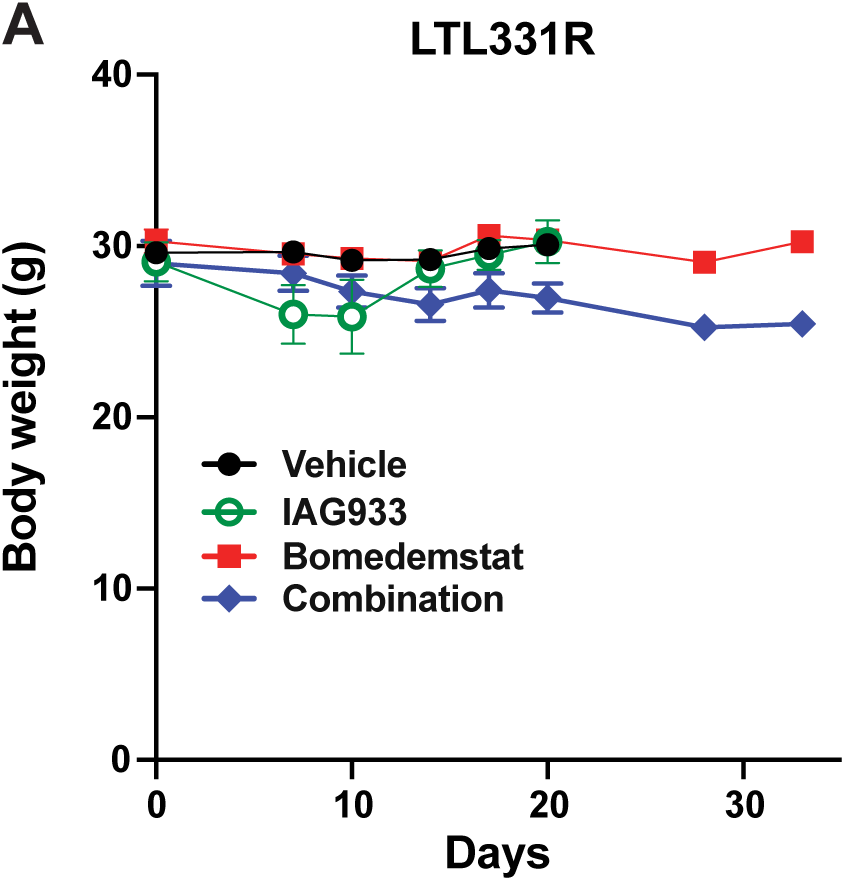
Body weights of mice treated with bomedemstat/IAG933 combination therapy. **A,** Body weights recorded for NSG mice bearing LTL331R PDX treated with bomedemstat monotherapy (dosed PO daily at 40 mg/kg), IAG933 (dosed PO daily at 240mg/kg), bomedemstat/IAG933 combination therapy with respective doses, or vehicle control (n=4-6/group). Days on treatment shown.

TEAD signatures.

**Table S1. TEAD signatures**
| **TEAD 73genes** | **TEAD 4genes** | **TEAD 6genes** | **TEAD 15genes** |
| --- | --- | --- | --- |
| CYR61 | CYR61 | CYR61 | CYR61 |
| AMOTL2 | AMOTL2 | AMOTL2 | AMOTL2 |
| ANXA3 | CTGF | ANXA3 | ANXA3 |
| MFAP5 | ANKRD1 | MFAP5 | MFAP5 |
| AXL |  | CTGF | AXL |
| ADRB2 |  | ANKRD1 | ADRB2 |
| DKK1 |  |  | DKK1 |
| FSTL1 |  |  | FSTL1 |
| SNAPC1 |  |  | SNAPC1 |
| PHLDB2 |  |  | PHLDB2 |
| F3 |  |  | F3 |
| CTGF |  |  | CTGF |
| AJUBA |  |  | AJUBA |
| ANKRD1 |  |  | ANKRD1 |
| MFAP5 |  |  | MFAP5 |
| EDN1 |  |  |  |
| FST |  |  |  |
| FOSL1 |  |  |  |
| DLC1 |  |  |  |
| TGM2 |  |  |  |
| FRMD6 |  |  |  |
| FAM46B |  |  |  |
| MAFF |  |  |  |
| CRIM1 |  |  |  |
| TBC1D2 |  |  |  |
| FGF2 |  |  |  |
| EPHA2 |  |  |  |
| TGFB2 |  |  |  |
| SERPINE1 |  |  |  |
| USCG |  |  |  |
| NOG |  |  |  |
| HBEGF |  |  |  |
| SAMD4A |  |  |  |
| TNFRSF12A |  |  |  |
| ITGB2 |  |  |  |
| SLFN12 |  |  |  |
| RND3 |  |  |  |
| LATS2 |  |  |  |
| APAP1L1 |  |  |  |
| TMEM45A |  |  |  |
| TUBB6 |  |  |  |
| PRICKLE1 |  |  |  |
| SFN |  |  |  |
| LAYN |  |  |  |
| CDC42EP3 |  |  |  |
| CCDC80 |  |  |  |
| SRGN |  |  |  |
| RBMS2 |  |  |  |
| ERBB2IP |  |  |  |
| KLF5 |  |  |  |
| PTRF |  |  |  |
| CCND1 |  |  |  |
| ACTN1 |  |  |  |
| HYI |  |  |  |
| GADD45B |  |  |  |
| ENC1 |  |  |  |
| TRIP6 |  |  |  |
| COL12A1 |  |  |  |
| ETS1 |  |  |  |
| ASAP1 |  |  |  |
| C10orf54 |  |  |  |
| EGFR |  |  |  |
| C1orf116 |  |  |  |
| ST6GALNAC5 |  |  |  |
| LIMA1 |  |  |  |
| LYPD6B |  |  |  |
| IFFO2 |  |  |  |
| PLEKHA7 |  |  |  |
| NFKBIZ |  |  |  |
| GAB2 |  |  |  |
| BICC1 |  |  |  |
| VANGL1 |  |  |  |
| PRKCH |  |  |  |

## References

1. Kratzer TB, Mazzitelli N, Star J, Dahut WL, Jemal A, Siegel RL. Prostate cancer statistics, 2025. CA: A Cancer Journal for Clinicians. 2025;75(6):485–97. doi:10.3322/caac.70028

2. Yamada Y, Beltran H. Clinical and Biological Features of Neuroendocrine Prostate Cancer. Curr Oncol Rep. 2021 Jan 12;23(2):15. doi:10.1007/s11912-020-01003-9

3. Gagnon R, Kish EK, Cook S, Takemura K, Cheng BYC, Bressler K, et al. Real-world Clinical Outcomes and Prognostic Factors in Neuroendocrine Prostate Cancer. Clinical Genitourinary Cancer. 2025 Feb 1;23(1):102274. doi:10.1016/j.clgc.2024.102274

4. Chen J, Shi M, Chuen Choi SY, Wang Y, Lin D, Zeng H, et al. Genomic alterations in neuroendocrine prostate cancer: A systematic review and meta-analysis. BJUI Compass. 2023;4(3):256–65. doi:10.1002/bco2.212

5. Ku SY, Rosario S, Wang Y, Mu P, Seshadri M, Goodrich ZW, et al. Rb1 and Trp53 cooperate to suppress prostate cancer lineage plasticity, metastasis, and antiandrogen resistance. Science. 2017 Jan 6;355(6320):78–83. doi:10.1126/science.aah4199

6. Chen CC, Tran W, Song K, Sugimoto T, Obusan MB, Wang L, et al. Temporal evolution reveals bifurcated lineages in aggressive neuroendocrine small cell prostate cancer trans-differentiation. Cancer Cell. 2023 Dec 11;41(12):2066–2082.e9. doi:10.1016/j.ccell.2023.10.009 PubMed PMID: 37995683.

7. Baine MK, Hsieh MS, Lai WV, Egger JV, Jungbluth AA, Daneshbod Y, et al. SCLC Subtypes Defined by ASCL1, NEUROD1, POU2F3, and YAP1: A Comprehensive Immunohistochemical and Histopathologic Characterization. J Thorac Oncol. 2020 Dec;15(12):1823–35. doi:10.1016/j.jtho.2020.09.009 PubMed PMID: 33011388; PubMed Central PMCID: PMC8362797.

8. Beltran H, Prandi D, Mosquera JM, Benelli M, Puca L, Cyrta J, et al. Divergent clonal evolution of castration-resistant neuroendocrine prostate cancer. Nat Med. 2016 Mar;22(3):298–305. doi:10.1038/nm.4045 PubMed PMID: 26855148; PubMed Central PMCID: PMC4777652.

9. Kumaraswamy A, Duan Z, Flores D, Zhang C, Sehrawat A, Hu YM, et al. LSD1 promotes prostate cancer reprogramming by repressing TP53 signaling independently of its demethylase function. JCI Insight. 2023 Aug 8;8(15):e167440. doi:10.1172/jci.insight.167440 PubMed PMID: 37440313; PubMed Central PMCID: PMC10445684.

10. Liu HM, Zhou Y, Chen HX, Wu JW, Ji SK, Shen L, et al. LSD1 in drug discovery: From biological function to clinical application. Medicinal Research Reviews. 2024;44(2):833– 66. doi:10.1002/med.22000

11. Han H, Wang Y, Curto J, Gurrapu S, Laudato S, Rumandla A, et al. Mesenchymal and stem-like prostate cancer linked to therapy-induced lineage plasticity and metastasis. Cell Reports. 2022 Apr;39(1):110595. doi:10.1016/j.celrep.2022.110595

12. Asrani K, Torres AF, Woo J, Vidotto T, Tsai HK, Luo J, et al. Reciprocal YAP1 loss and INSM1 expression in neuroendocrine prostate cancer. The Journal of Pathology. 2021;255(4):425–37. doi:10.1002/path.5781

13. Abida W, Cyrta J, Heller G, Prandi D, Armenia J, Coleman I, et al. Genomic correlates of clinical outcome in advanced prostate cancer. Proceedings of the National Academy of Sciences. 2019 Jun 4;116(23):11428–36. doi:10.1073/pnas.1902651116

14. Sun C, Shi Y, Xu LL, Nageswararao C, Davis LD, Segawa T, et al. Androgen receptor mutation (T877A) promotes prostate cancer cell growth and cell survival. Oncogene. 2006 Jun;25(28):3905–13. doi:10.1038/sj.onc.1209424

15. Cunningham D, You Z. In vitro and in vivo model systems used in prostate cancer research. J Biol Methods. 2015;2(1):e17. doi:10.14440/jbm.2015.63 PubMed PMID: 26146646; PubMed Central PMCID: PMC4487886.

16. Haffner MC, Bhamidipati A, Tsai HK, Esopi DM, Vaghasia AM, Low JY, et al. PHENOTYPIC CHARACTERIZATION OF TWO NOVEL CELL LINE MODELS OF CASTRATION RESISTANT PROSTATE CANCER. Prostate. 2021 Nov;81(15):1159–71. doi:10.1002/pros.24210 PubMed PMID: 34402095; PubMed Central PMCID: PMC8460612.

17. Brennen WN, Zhu Y, Coleman IM, Dalrymple SL, Antony L, Patel RA, et al. Resistance to androgen receptor signaling inhibition does not necessitate development of neuroendocrine prostate cancer. JCI Insight. 6(8):e146827. doi:10.1172/jci.insight.146827 PubMed PMID: 33724955; PubMed Central PMCID: PMC8119192.

18. Han G, Buchanan G, Ittmann M, Harris JM, Yu X, DeMayo FJ, et al. Mutation of the androgen receptor causes oncogenic transformation of the prostate. Proceedings of the National Academy of Sciences. 2005 Jan 25;102(4):1151–6. doi:10.1073/pnas.0408925102

19. Zhu Y, Dalrymple SL, Coleman I, Zheng SL, Xu J, Hooper JE, et al. Role of androgen receptor splice variant-7 (AR-V7) in prostate cancer resistance to 2nd-generation androgen receptor signaling inhibitors. Oncogene. 2020 Nov;39(45):6935–49. doi:10.1038/s41388-020-01479-6

20. Han W, Liu M, Han D, Li M, Toure AA, Wang Z, et al. RB1 loss in castration-resistant prostate cancer confers vulnerability to LSD1 inhibition. Oncogene. 2022 Feb;41(6):6. doi:10.1038/s41388-021-02135-3

21. Sehrawat A, Gao L, Wang Y, Bankhead A, McWeeney SK, King CJ, et al. LSD1 activates a lethal prostate cancer gene network independently of its demethylase function. Proc Natl Acad Sci USA. 2018 May;115(18). doi:10.1073/pnas.1719168115

22. Augert A, Eastwood E, Ibrahim AH, Wu N, Grunblatt E, Basom R, et al. Targeting NOTCH activation in small cell lung cancer through LSD1 inhibition. Science Signaling. 2019 Feb 5;12(567):eaau2922. doi:10.1126/scisignal.aau2922

23. Nguyen EM, Taniguchi H, Chan JM, Zhan YA, Chen X, Qiu J, et al. Targeting Lysine-Specific Demethylase 1 Rescues Major Histocompatibility Complex Class I Antigen Presentation and Overcomes Programmed Death-Ligand 1 Blockade Resistance in SCLC. Journal of Thoracic Oncology. 2022 Aug 1;17(8):1014–31. doi:10.1016/j.jtho.2022.05.014

24. Romero R, Chu T, González Robles TJ, Smith P, Xie Y, Kaur H, et al. The neuroendocrine transition in prostate cancer is dynamic and dependent on ASCL1. Nat Cancer. 2024 Nov;5(11):1641–59. doi:10.1038/s43018-024-00838-6

25. Tang F, Xu D, Wang S, Wong CK, Martinez-Fundichely A, Lee CJ, et al. Chromatin profiles classify castration-resistant prostate cancers suggesting therapeutic targets. Science. 2022 May 27;376(6596):eabe1505. doi:10.1126/science.abe1505

26. Perillo B, Tramontano A, Pezone A, Migliaccio A. LSD1: more than demethylation of histone lysine residues. Exp Mol Med. 2020 Dec;52(12):12. doi:10.1038/s12276-020-00542-2

27. Estève PO, Chin HG, Benner J, Feehery GR, Samaranayake M, Horwitz GA, et al. Regulation of DNMT1 stability through SET7-mediated lysine methylation in mammalian cells. Proceedings of the National Academy of Sciences. 2009 Mar 31;106(13):5076–81. doi:10.1073/pnas.0810362106

28. Pearson JD, Huang K, Pacal M, McCurdy SR, Lu S, Aubry A, et al. Binary pan-cancer classes with distinct vulnerabilities defined by pro- or anti-cancer YAP/TEAD activity. Cancer Cell. 2021 Aug;39(8):1115–1134.e12. doi:10.1016/j.ccell.2021.06.016

29. Dumbrava EE, Shapiro GI, Parikh AR, Johnson ML, Tolcher AW, Thompson JA, et al. Phase 1 Study of Rezatapopt, a p53 Reactivator, in TP53 Y220C–Mutated Tumors. New England Journal of Medicine. 2026 Feb 25;394(9):872–83. doi:10.1056/NEJMoa2508820

30. Dumbrava EE, Stinchcombe TE, Gounder M, Cote GM, Hanna GJ, Sumrall B, et al. Milademetan in Advanced Solid Tumors with MDM2 Amplification and Wild-type TP53: Preclinical and Phase II Clinical Trial Results. Clin Cancer Res. 2025 Oct 15;31(20):4255– 64. doi:10.1158/1078-0432.CCR-25-0762

31. The Biology of YAP/TAZ: Hippo Signaling and Beyond | Physiological Reviews | American Physiological Society [Internet]. [cited 2026 Apr 1]. Available from: https://journals.physiology.org/doi/full/10.1152/physrev.00005.2014?r=1&fst=0&l=ri

32. Bata N, Chaikuad A, Bakas NA, Limpert AS, Lambert LJ, Sheffler DJ, et al. Inhibitors of the Hippo Pathway Kinases STK3/MST2 and STK4/MST1 Have Utility for the Treatment of Acute Myeloid Leukemia. J Med Chem. 2022 Jan 27;65(2):1352–69. doi:10.1021/acs.jmedchem.1c00804

33. Chapeau EA, Sansregret L, Galli GG, Chène P, Wartmann M, Mourikis TP, et al. Direct and selective pharmacological disruption of the YAP–TEAD interface by IAG933 inhibits Hippo-dependent and RAS–MAPK-altered cancers. Nat Cancer. 2024 Jul;5(7):1102–20. doi:10.1038/s43018-024-00754-9

34. Wang S, Yang C, Tang J, Wang K, Cheng H, Yao S, et al. LSD1 is a targetable vulnerability in gastric cancer harboring TP53 frameshift mutations. Clin Epigenet. 2025 Feb 18;17(1):26. doi:10.1186/s13148-025-01829-9

35. Lachowiez C, Kaempf A, Lehn C, Lundberg Williams T, Sheldon G, Leonard J, et al. Preliminary safety and efficacy results of A phase ib investigation of the LSD1 inhibitor iadademstat (ORY-1001) in combination with azacitidine and venetoclax in newly-diagnosed AML. Blood. 2025 Nov 3;146(Supplement 1):1649. doi:10.1182/blood-2025-1649

36. Bauer TM, Besse B, Martinez-Marti A, Trigo JM, Moreno V, Garrido P, et al. Phase I, Open-Label, Dose-Escalation Study of the Safety, Pharmacokinetics, Pharmacodynamics, and Efficacy of GSK2879552 in Relapsed/Refractory SCLC. Journal of Thoracic Oncology. 2019 Oct 1;14(10):1828–38. doi:10.1016/j.jtho.2019.06.021

37. Montalban-Bravo G, DiNardo CD, Short N, Kadia TM, Ravandi F, Chien KS, et al. A Phase I/II Study of Seclidemstat, an LSD1 Inhibitor, and Azacitidine for Patients with Myelodysplastic Syndromes and Chronic Myelomonocytic Leukemia. Blood. 2022 Nov 15;140(Supplement 1):9771–2. doi:10.1182/blood-2022-170138

38. Nouruzi S, Ganguli D, Tabrizian N, Kobelev M, Sivak O, Namekawa T, et al. ASCL1 activates neuronal stem cell-like lineage programming through remodeling of the chromatin landscape in prostate cancer. Nat Commun. 2022 Apr 27;13(1):1. doi:10.1038/s41467-022-29963-5

39. Hata K, Okano M, Lei H, Li E. Dnmt3L cooperates with the Dnmt3 family of de novo DNA methyltransferases to establish maternal imprints in mice. Development. 2002 Apr;129(8):1983–93. doi:10.1242/dev.129.8.1983 PubMed PMID: 11934864.

40. Okano M, Bell DW, Haber DA, Li E. DNA methyltransferases Dnmt3a and Dnmt3b are essential for de novo methylation and mammalian development. Cell. 1999 Oct 29;99(3):247–57. doi:10.1016/s0092-8674(00)81656-6 PubMed PMID: 10555141.

41. Chen T, Ueda Y, Dodge JE, Wang Z, Li E. Establishment and maintenance of genomic methylation patterns in mouse embryonic stem cells by Dnmt3a and Dnmt3b. Mol Cell Biol. 2003 Aug;23(16):5594–605. doi:10.1128/MCB.23.16.5594-5605.2003 PubMed PMID: 12897133; PubMed Central PMCID: PMC166327.

42. Tahiliani M, Koh KP, Shen Y, Pastor WA, Bandukwala H, Brudno Y, et al. Conversion of 5-methylcytosine to 5-hydroxymethylcytosine in mammalian DNA by MLL partner TET1. Science. 2009 May 15;324(5929):930–5. doi:10.1126/science.1170116 PubMed PMID: 19372391; PubMed Central PMCID: PMC2715015.

43. Cejas P, Xie Y, Font-Tello A, Lim K, Syamala S, Qiu X, et al. Subtype heterogeneity and epigenetic convergence in neuroendocrine prostate cancer. Nat Commun. 2021 Oct 1;12(1):1. doi:10.1038/s41467-021-26042-z

44. Zanconato F, Forcato M, Battilana G, Azzolin L, Quaranta E, Bodega B, et al. Genome-wide association between YAP/TAZ/TEAD and AP-1 at enhancers drives oncogenic growth. Nat Cell Biol. 2015 Sep;17(9):1218–27. doi:10.1038/ncb3216

45. Pflug BR, Reiter RE, Nelson JB. Caveolin expression is decreased following androgen deprivation in human prostate cancer cell lines. The Prostate. 1999;40(4):269–73. doi:10.1002/(SICI)1097-0045(19990901)40:4<269::AID-PROS9>3.0.CO;2-6

46. Mertz KD, Setlur SR, Dhanasekaran SM, Demichelis F, Perner S, Tomlins S, et al. Molecular Characterization of *TMPRSS2-ERG* Gene Fusion in the NCI-H660 Prostate Cancer Cell Line: A New Perspective for an Old Model. Neoplasia. 2007 Mar 1;9(3):200-IN3. doi:10.1593/neo.07103

47. van Bokhoven A, Varella-Garcia M, Korch C, Johannes WU, Smith EE, Miller HL, et al. Molecular characterization of human prostate carcinoma cell lines. The Prostate. 2003;57(3):205–25. doi:10.1002/pros.10290

48. Lin D, Wyatt AW, Xue H, Wang Y, Dong X, Haegert A, et al. High Fidelity Patient-Derived Xenografts for Accelerating Prostate Cancer Discovery and Drug Development. Cancer Research. 2014 Feb 15;74(4):1272–83. doi:10.1158/0008-5472.CAN-13-2921-T

49. Yu X, Zhan X, D’Costa J, Tanavde VM, Ye Z, Peng T, et al. Lentiviral vectors with two independent internal promoters transfer high-level expression of multiple transgenes to human hematopoietic stem-progenitor cells. Mol Ther. 2003 Jun;7(6):827–38. doi:10.1016/s1525-0016(03)00104-7 PubMed PMID: 12788657.

50. Moffat J, Grueneberg DA, Yang X, Kim SY, Kloepfer AM, Hinkle G, et al. A lentiviral RNAi library for human and mouse genes applied to an arrayed viral high-content screen. Cell. 2006 Mar 24;124(6):1283–98. doi:10.1016/j.cell.2006.01.040 PubMed PMID: 16564017.

51. Junqueira Alves C, Dariolli R, Haydak J, Kang S, Hannah T, Wiener RJ, et al. Plexin-B2 orchestrates collective stem cell dynamics via actomyosin contractility, cytoskeletal tension and adhesion. Nat Commun. 2021 Oct 14;12(1):6019. doi:10.1038/s41467-021-26296-7 PubMed PMID: 34650052; PubMed Central PMCID: PMC8517024.

52. Brennen WN, Rosen DM, Wang H, Isaacs JT, Denmeade SR. Targeting Carcinoma-Associated Fibroblasts Within the Tumor Stroma With a Fibroblast Activation Protein-Activated Prodrug. J Natl Cancer Inst. 2012 Sep 5;104(17):1320–34. doi:10.1093/jnci/djs336

53. Bankhead P, Loughrey MB, Fernández JA, Dombrowski Y, McArt DG, Dunne PD, et al. QuPath: Open source software for digital pathology image analysis. Sci Rep. 2017 Dec 4;7(1):16878. doi:10.1038/s41598-017-17204-5

54. Dobin A, Davis CA, Schlesinger F, Drenkow J, Zaleski C, Jha S, et al. STAR: ultrafast universal RNA-seq aligner. Bioinformatics. 2013 Jan 1;29(1):15–21. doi:10.1093/bioinformatics/bts635

55. XenofilteR: computational deconvolution of mouse and human reads in tumor xenograft sequence data | BMC Bioinformatics | Springer Nature Link [Internet]. [cited 2026 May 15]. Available from: https://link.springer.com/article/10.1186/s12859-018-2353-5

56. Lawrence M, Huber W, Pagès H, Aboyoun P, Carlson M, Gentleman R, et al. Software for Computing and Annotating Genomic Ranges. PLOS Computational Biology. 2013 Aug 8;9(8):e1003118. doi:10.1371/journal.pcbi.1003118

57. Ritchie ME, Phipson B, Wu D, Hu Y, Law CW, Shi W, et al. limma powers differential expression analyses for RNA-sequencing and microarray studies. Nucleic Acids Res. 2015 Apr 20;43(7):e47. doi:10.1093/nar/gkv007

58. Subramanian A, Tamayo P, Mootha VK, Mukherjee S, Ebert BL, Gillette MA, et al. Gene set enrichment analysis: A knowledge-based approach for interpreting genome-wide expression profiles. Proceedings of the National Academy of Sciences. 2005 Oct 25;102(43):15545–50. doi:10.1073/pnas.0506580102

59. Hänzelmann S, Castelo R, Guinney J. GSVA: gene set variation analysis for microarray and RNA-Seq data. BMC Bioinformatics. 2013 Jan 16;14(1):7. doi:10.1186/1471-2105-14-7

60. Yegnasubramanian S, Lin X, Haffner MC, DeMarzo AM, Nelson WG. Combination of methylated-DNA precipitation and methylation-sensitive restriction enzymes (COMPARE-MS) for the rapid, sensitive and quantitative detection of DNA methylation. Nucleic Acids Res. 2006 Feb 1;34(3):e19. doi:10.1093/nar/gnj022

61. Anders NM, Liu J, Wanjiku T, Giovinazzo H, Zhou J, Vaghasia A, et al. Simultaneous quantitative determination of 5-aza-2’-deoxycytidine genomic incorporation and DNA demethylation by liquid chromatography tandem mass spectrometry as exposure-response measures of nucleoside analog DNA methyltransferase inhibitors. J Chromatogr B Analyt Technol Biomed Life Sci. 2016 Jun 1;1022:38–45. doi:10.1016/j.jchromb.2016.03.029 PubMed PMID: 27082761; PubMed Central PMCID: PMC4866886.

62. Cerami E, Gao J, Dogrusoz U, Gross BE, Sumer SO, Aksoy BA, et al. The cBio Cancer Genomics Portal: An Open Platform for Exploring Multidimensional Cancer Genomics Data. Cancer Discov. 2012 May 9;2(5):401–4. doi:10.1158/2159-8290.CD-12-0095

63. Zhang X, Coleman IM, Brown LG, True LD, Kollath L, Lucas JM, et al. SRRM4 Expression and the Loss of REST Activity May Promote the Emergence of the Neuroendocrine Phenotype in Castration-Resistant Prostate Cancer. Clin Cancer Res. 2015 Oct 15;21(20):4698–708. doi:10.1158/1078-0432.CCR-15-0157 PubMed PMID: 26071481; PubMed Central PMCID: PMC4609255.

64. Lapuk AV, Wu C, Wyatt AW, McPherson A, McConeghy BJ, Brahmbhatt S, et al. From sequence to molecular pathology, and a mechanism driving the neuroendocrine phenotype in prostate cancer. The Journal of Pathology. 2012;227(3):286–97. doi:10.1002/path.4047

65. Ertel A, Dean JL, Rui H, Liu C, Witkiewicz A, Knudsen KE, et al. RB-pathway disruption in breast cancer: Differential association with disease subtypes, disease-specific prognosis and therapeutic response. Cell Cycle. 2010 Oct 15;9(20):4153–63. doi:10.4161/cc.9.20.13454 PubMed PMID: 20948315.

66. Tsai HK, Lehrer J, Alshalalfa M, Erho N, Davicioni E, Lotan TL. Gene expression signatures of neuroendocrine prostate cancer and primary small cell prostatic carcinoma. BMC Cancer. 2017 Nov 13;17(1):759. doi:10.1186/s12885-017-3729-z

67. Bluemn EG, Coleman IM, Lucas JM, Coleman RT, Hernandez-Lopez S, Tharakan R, et al. Androgen Receptor Pathway-Independent Prostate Cancer Is Sustained through FGF Signaling. Cancer Cell. 2017 Oct;32(4):474–489.e6. doi:10.1016/j.ccell.2017.09.003

68. Nguyen LT, Tretiakova MS, Silvis MR, Lucas J, Klezovitch O, Coleman I, et al. ERG Activates the YAP1 Transcriptional Program and Induces the Development of Age-Related Prostate Tumors. Cancer Cell. 2015 Jun 8;27(6):797–808. doi:10.1016/j.ccell.2015.05.005 PubMed PMID: 26058078.

69. Wang Y, Xu X, Maglic D, Dill MT, Mojumdar K, Ng PKS, et al. Comprehensive Molecular Characterization of the Hippo Signaling Pathway in Cancer. Cell Reports. 2018 Oct 30;25(5):1304–1317.e5. doi:10.1016/j.celrep.2018.10.001 PubMed PMID: 30380420.

